# Modulation of Alzheimer’s Disease Brain Pathology in Mice by Gut Bacterial Deletion: The Role of Il-17a and Microglial MyD88

**DOI:** 10.1101/2023.12.06.570400

**Authors:** Wenlin Hao, Qinghua Luo, Inge Tomic, Wenqiang Quan, Tobias Hartmann, Michael D. Menger, Klaus Fassbender, Yang Liu

## Abstract

Gut bacteria regulate brain pathology of Alzheimer’s disease (AD) patients and animal models; however, the underlying mechanism remains unclear. In this study, 3-month-old APP-transgenic female mice with and without knock-out of *Il-17a* gene, or haploinsufficiency of MyD88 in microglia were treated with antibiotics-supplemented or normal drinking water for 2 months. Antibiotic treatment eradicated gut bacteria, particularly in the phyla *Bacteroidetes* and *Firmicutes*, and reduced Il-17a-expressing CD4-positive T lymphocytes. Deletion of gut bacteria inhibited inflammatory activation in the brain and microglia, and reduced cerebral Aβ levels in APP-transgenic mice, which was abolished by deficiency of Il-17a or haploinsufficiency of MyD88 in microglia. As possible mechanisms regulating Aβ pathology, deletion of gut bacteria inhibited β-secretase activity and increased the expression of Abcb1 and Lrp1 in the brain or at the blood-brain barrier, which were also reversed by the absence of Il-17a. Interestingly, a crossbreeding experiment between APP-transgenic mice and *Il-17a* knockout mice further showed that deficiency of Il-17a had already increased Abcb1 and Lrp1 expression at the blood-brain barrier. Thus, deletion of gut bacteria attenuates inflammatory activation and amyloid pathology in APP-transgenic mice via Il-17a and microglial MyD88-involved signalling pathways. Our study contributes to a better understanding of the gut-brain axis in AD pathophysiology.

## Introduction

Changes in bacterial components have been observed in the gut of Alzheimer’s disease (AD) patients and mouse models, although the characteristic phenotype of bacteria has not yet been determined (Vogt et al. 2017; Dunham et al. 2022; Meyer et al. 2022; Zhang T et al. 2023). In a prospective study of cognitively healthy individuals, a decrease in butyrate-producing bacterial species (e.g., *Roseburia inulinivorans* and *R. faecis*) or an increase in pro-inflammatory bacteria (e.g., *Veillonella dispar* and *V. atypica*) was associated with subjective cognitive decline during a follow-up period of 2-4 years (Ma et al. 2023). Transplantation of gut bacteria from AD patients to gut bacteria-depleted rats leads to impaired neurogenesis and cognitive deficits (Grabrucker et al. 2023). Gut bacteria-free Alzheimer’s precursor protein (APP)-transgenic or *App^NL-G-F^* knock-in mice show a reduction in cerebral amyloid β peptide (Aβ) deposition and microgliosis (Minter et al. 2016; Harach et al. 2017; Minter et al. 2017; Dodiya et al. 2019; Kaur et al. 2021). Therefore, gut bacteria contribute to AD pathogenesis; however, how gut bacteria regulate microglial activation and Aβ pathology remains unclear.

Intestinal bacteria-derived short-chain free fatty acids (SCFAs; i.e., acetate, butyrate, and propionate) have been shown to promote microglial maturation, innate immune responses, and energy metabolism in the homeostatic state (Erny et al. 2015; Erny et al. 2021). Administration of SCFAs to germ-free or antibiotic-treated AD mice restores microglial proliferation and inflammatory activation; however, the effects on Aβ phagocytosis and Aβ accumulation in the brain are inconsistent across different studies (Colombo et al. 2021; Erny et al. 2021; Xie et al. 2023). Whether SCFAs act directly on microglia remains unclear. We found that SCFA receptors, G protein-coupled receptor (Gpr) 41 and Gpr43 are absent in murine microglia (Quan et al. 2021); however, deficiency of Gpr41 and Gpr43 likely inhibits microglial maturation under physiological conditions (Erny et al. 2015) and increases microglial density and Aβ deposits in the brain of APP-transgenic mice (Zhou et al. 2023). Deficiency of the butyrate receptor Gpr109a, which is highly expressed in microglia (Quan et al. 2021), has limited effects on microglial activation (Zhou et al. 2023).

It is known that intestinal bacteria regulate T lymphocytes, which potentially regulate brain cells. Germ-free mice generate fewer interleukin (Il)-17a-producing CD4(+) T help (Th17) lymphocytes but more CD4(+)CD25(+)Foxp3(+) regulatory T (Treg) cells in the gut and spinal cord, which is associated with resistance to experimental autoimmune encephalomyelitis in mice (Lee et al. 2011). Deletion of intestinal bacteria by antibiotics reduces the accumulation of Il-17a-producing γδ T cells in the leptomeninges and ameliorates brain injury in a stroke mouse model (Benakis et al. 2016). In APP-transgenic mice, reductions in cerebral Aβ deposition and microglia after gut antibiotic treatments correlate with increased levels of Foxp3+ Treg cells in blood and brain (Minter et al. 2017). However, transient depletion of Treg cells was shown to regulate microglia or/and infiltrated macrophages with differential effects on Aβ clearance and cognitive protection in two studies (Baruch et al. 2015; Dansokho et al. 2016). Our recent study showed that deficiency of p38α-MAPK in peripheral myeloid cells decreases Th17 cells, which possibly increases microglial activation and Aβ clearance in AD mice (Luo et al. 2022). It is yet unknown whether Il-17a-expressing cells mediate the effects of gut bacteria on microglia.

Intestinal bacteria may also release structural components into the blood and directly activate microglia. Blood concentrations of lipopolysaccharide (LPS) together with inflammatory cytokines, e.g. interleukin (IL)-1β and tumor necrosis factor (TNF)-α, are increased in AD patients compared to non-AD individuals with and without cognitive impairment (Marizzoni et al. 2023). Components of *Porphyromonas gingivalis*, a bacterium often existing in chronic periodontitis, were found in the brains of AD patients (Dominy et al. 2019). We have shown that deficiency of MyD88, an adaptor protein downstream of most Toll-like receptors (O’Neill et al. 2013), regulates microglial activation and cerebral Aβ pathology in APP-transgenic mice (Hao et al. 2011; Quan et al. 2021). It will be interesting to learn whether the innate immune signaling mediates the effects of intestinal bacteria on AD pathogenesis.

In this study, we treated female APP-transgenic mice with and without an antibiotic cocktail in the drinking water and observed that deletion of intestinal bacteria reduced inflammatory activation and Aβ load in the brain. These effects could be abolished by general knockout of *Il-17a* gene or conditional knockout of *Myd88* gene in microglia.

## Materials and methods

### Animal models and cross-breeding

APP/PS1-transgenic mice (APP^tg^) over-expressing human mutated APP (KM670/671NL) and PS1 (L166P) under Thy-1 promoters (Radde et al. 2006) were gifts from M. Jucker, Hertie Institute for Clinical Brain Research, Tübingen, Germany. *Il-17a* knockout (Il17a^-/-^) mice were kindly provided by Y. Iwakura, Tokyo University of Science, Japan (Nakae et al. 2002). Il-17a-deficient AD mice were created by cross-breeding APP/PS1-transgenic mice and Il17a^-/-^ mice to obtain APP^tg^/Il17a^-/-^ genotype in our previous study (Luo et al. 2022). Microglial MyD88-haplodefficient AD mice with the APP^tg^MyD88^fl/wt^Cre^+/−^ genotype were generated in our previous study (Quan et al. 2021) by mating APP/PS1-transgenic mice with *Myd88*-floxed mice (Hou et al. 2008) and Cx3Cr1-CreERT2 mice (Goldmann et al. 2013) and intraperitoneal injection of tamoxifen (100 mg/kg; Sigma-Aldrich Chemie, Taufkirchen, Germany) once daily for 5 days at 1 month of age. APP/PS1-transgenic mice were also cross-bred with Il-17a-eGFP reporter mice (Il17a^GFP/GFP^; kindly provided by R. Flavell, Yale University, USA) to get APP^tg^/Il17a^GFP/wt^ of genotype, in which eGFP is expressed under the control of endogenous *Il-17a* gene promoter (Esplugues et al. 2011). All mice used in this project were on C57BL/6 genetic background.

### Deletion of intestinal bacteria with antibiotics in drinking water

Various AD mice at 3 months of age, and 3 or 22-month-old C57BL/6J mice were treated with vancomycin (500mg/L), ampicillin (1g/L), neomycin sulfate (1g/L), streptomycin (1g/L), and metronidazole (1g/L) (all antibiotics were bought from Sigma-Aldrich Chemie GmbH, Taufkirchen, Germany) in the drinking water for 2 months to delete intestinal bacteria according to a published protocol (Liu S et al. 2016). The water with antibiotics was changed every 7 days. Animal experiments were conducted in accordance with national rules and ARRIVE guidelines, and authorized by Landesamt für Verbraucherschutz, Saarland, Germany (registration numbers: 56/2015, 46/2017 and 34/2019).

### Tissue collection and isolation of blood vessels

Animals were euthanized at 5 months of age by inhalation of overdose isoflurane and perfused with ice-cold PBS. The brain was removed and divided. The left hemisphere was immediately fixed in 4% paraformaldehyde (Sigma-Aldrich) in PBS and embedded in paraffin for immunohistochemistry. A 0.5-mm-thick piece of tissue was sagittally cut from the right hemisphere. The cortex and hippocampus were carefully separated and homogenized in TRIzol (Thermo Fisher Scientific, Darmstadt, Germany) for RNA isolation. The remainder of the right hemisphere was snap-frozen in liquid nitrogen and stored at -80°C until biochemical analysis. The appendix together with a 0.5-cm-long segment of the neighboring colon was also collected, snap-frozen and stored at -80°C for isolation of intestinal bacteria.

To isolate brain blood vessels, the cortex and hippocampus from 5-month-old APP-transgenic mice were carefully dissected and brain vessel fragments were isolated as we did previously (Quan et al. 2021). Briefly, brain tissues were homogenized in HEPES-contained Hanks’ balanced salt solution (HBSS) and centrifuged at 4,400g in HEPES-HBSS buffer supplemented with dextran from *Leuconostoc spp*. (molecular weight ∼70,000; Sigma-Aldrich) to delete myelin. The vessel pellet was re-suspended in HEPES-HBSS buffer supplemented with 1% bovine serum albumin (Sigma-Aldrich) and filtered with 20 μm-mesh. The blood vessel fragments were collected on the top of filter and stored at -80°C for biochemical analysis.

### Intestinal bacterial collection and 16S rRNA sequencing

Bacterial DNA was extracted from intestinal bacteria in the frozen appendix and colon with QIAamp Fast DNA Stool Mini Kit (Qiagen, Hilden, Germany). The V3 -V4 region of the 16S rRNA-encoding gene was amplified with the barcode fusion primers (338F: 5-ACTCCTACGGGAGGCAGCAG-3, and 806R: 5-GGACTACHVGGGTWTCTAAT-3). After purification, PCR products were used for constructing libraries and sequenced on the Illumina MiSeq platform at Majorbio Co. Ltd. (Shanghai, China). The raw data was processed on Qiime2 (https://qiime2.org/), reducing sequencing and PCR errors, and denoising to get the operational taxonomic unit (OTU) consensus sequences, which were mapped to the 16S Mothur-Silva SEED r119 database (http://www.mothur.org/). Alpha diversity including Sobs, Shannon, Ace, Chao and Simpson indexes were used for the analysis of bacterial richness and diversity in a single mouse. Principal coordinate analysis (PCoA) and analysis of similarity (ANOSIM) were used for β-diversity analysis to compare bacterial compositions on genus level between APP-transgenic mice with and without antibiotic treatments. The difference of bacterial compositions on phylum level between these two groups were also compared with Wilcoxon rink sum test. All the analysis was performed using cloud-based tools with default analysis parameters (https://cloud.majorbio.com/page/tools.html).

### Positive selection of CD11b-positive and CD4-positive cells from the brain and spleen, respectively

The brain tissue (hippocampus and cortex) and spleen of 5-month-old APP-transgenic mice with and without treatments with antibiotics were prepared for single-cell suspensions using Neural Tissue Dissociation Kit (papain-based) and Spleen Dissociation Kit (mouse), respectively (Miltenyi Biotec GmbH, Bergisch Gladbach, Germany). After blocking with 50 µg/ml CD16/CD32 antibody (clone 2.4G2; BioXCell, Lebanon, USA), CD11b-positive brain cells were selected from the brain with microbeads-conjugated CD11b antibody (clone M1/70.15.11.5; Miltenyi Biotec) and CD4-positive spleen cells from the spleen with Dynabeads magnetic beads-conjugated CD4 antibody (clone L3T4; Thermo Fisher Scientific). Lysis buffer was immediately added to selected cells and total RNA was isolated using RNeasy Plus Mini Kit (Qiagen).

### Microglial A**β** phagocytosis assay

Five-month-old APP-transgenic mice received an intraperitoneal injection of 10 mg/kg methoxy-XO4 (Bio-Techne GmbH, Wiesbaden-Nordenstadt, Germany) (2 mg/ml in a 1:1 mixture of DMSO and 0.9% NaCl [pH 12] (v/v)) after treatment with and without antibiotics according to a published protocol (Lau et al. 2021). Methoxy-XO4 binds to β-sheet secondary structure of Aβ aggregates. Three hours later, a single cell suspension from the hippocampus and cortex was prepared using Neural Tissue Dissociation Kit (papain-based) (Miltenyi Biotec GmbH). After blocking with CD16/CD32 antibody (clone 2.4G2; BioXCell) and subsequent staining with PE-Cy5-conjugated CD11b antibody (clone M1/70; Thermo Fisher Scientific), fibrillar Aβ-containing CD11b-positive brain cells were detected by BD FACSVerse™ flow cytometry (BD Biosciences; Heidelberg, Germany).

### Flow cytometric detection of Il17a-eGFP reporter in intestinal cells

A published protocol (Couter and Surana 2016) was used to prepare single cell suspensions from both lamina propria and Peyer’s patches of the small intestine of 5-month-old APP^tg^/Il17a^GFP/wt^ mice with and without two months of antibiotic treatment. After staining with APC-conjugated rat anti-CD4 monoclonal antibody (clone: GK1.5; Thermo Fisher Scientific), eGFP-expressing CD4-positive cells were detected by BD FACSCanto™ II flow cytometry (BD Biosciences).

### Histological analysis

Serial 50-μm-thick sagittal sections were cut from the paraffin-embedded hemisphere. Four neighboring sections with 300 µm of interval were deparaffinized, labeled with rabbit anti-ionized calcium-binding adapter molecule (Iba)-1 antibody (Wako Chemicals, Neuss, Germany) and VectaStain *Elite* ABC-HRP kit (Cat.-No.: PK-6100, Vector Laboratories, Burlingame, USA), and visualized with diaminobenzidine (Sigma-Aldrich). Iba-1-positive microglia/brain macrophages were counted with Optical Fractionator in the hippocampus on a Zeiss AxioImager.Z2 microscope (Carl Zeiss Microscopy GmbH, Göttingen, Germany) equipped with a Stereo Investigator system (MBF Bioscience, Williston), as we did previously (Liu Y et al. 2014).

To evaluate the cerebral Aβ level, after deparaffinization, 4 serial brain sections from each animal were stained with rabbit anti-human Aβ antibody (clone D12B2; Cell Signaling Technology Europe, Frankfurt am Main, Germany) and Cy3-conjuagted goat anti-rabbit IgG (Jackson ImmunoResearch Europe Ltd. Cambridge, UK), or with methoxy-XO4 (Bio-Techne GmbH). After mounting, the whole section including hippocampus and cortex was imaged with Microlucida (MBF Bioscience) and merged. The positive staining and brain region analyzed were measured for the area with Image J tool “Analyse Particles” (https://imagej.nih.gov/ij/). The threshold for all compared samples was manually set and kept constantly. The percentage of Aβ coverage in the brain was calculated.

### Analysis of microglial morphology

For the analysis of microglial morphology, our established protocol and Fiji Image J were used (Luo et al. 2022). Paraffin-embedded 50-µm sagittal brain sections were used as described above. After fluorescent co-staining with Iba-1 and Aβ antibodies, total 10 Aβ plaques/mouse were randomly selected from the cortex dorsal to hippocampus and imaged under 40× objective with Z-stack scanning with 1 µm of interval. The serial images were Z-projected with maximal intensity, 8-bit grayscale transformed, Unsharp-Mask filter and despeckle-treated, and binarized to obtain a black and white image. The cells with complete nucleus and branches and without overlapping with neighboring cells were chosen for analysis. The single-pixel background noise was eliminated and the gaps along processes were filled under the view of the original image of the cell. The processed image was skeletonized and analyzed with the plugin Analyze Skeleton (2D/3D) (http://imagej.net/AnalyzeSkeleton) for the total number of primary branches, length of all branches, and the number of branch endpoints of each microglia. The whole analysis was done blinded to genotypes.

### Western blot analysis

Frozen brain tissues and blood vessel isolates were homogenized in RIPA buffer (50mM Tris [pH 8.0], 150mM NaCl, 0.1% SDS, 0.5% sodiumdeoxy-cholate, 1% NP-40, and 5mM EDTA) supplemented with protease inhibitor cocktail (Roche Applied Science, Mannheim, Germany) on ice. The proteins were separated by 10% or 12% Tris-glycine SDS/PAGE. Before loading on PAGE gel, vessel preparations were sonicated. Western blots were performed using rabbit monoclonal antibody against Abcb1 (clone E1Y7S) and rabbit polyclonal antibody against Lrp1 (Cat.-No.: 64099) (both antibodies were bought from Cell Signaling Technology), as well as rabbit polyclonal antibody against claudin-5 (Cat.-No.: GTX49370; GeneTex, Hsinchu, China). The detected proteins were visualized via the Plus-ECL method (PerkinElmer, Waltham, USA). To quantify proteins of interest, rabbit monoclonal antibody against β-actin (clone 13E5) or rabbit polyclonal antibody against α-tubulin (Cat.-No.: 2144) (both antibodies were from Cell Signaling Technology) was used as a protein loading control. Densitometric analysis of band densities was performed with Image-Pro Plus software version 6.0.0.260 (Media Cybernetics, Rockville, MD). For each sample, the protein level was calculated as a ratio of target protein/loading control per sample.

### Brain homogenates and Aβ ELISA

The frozen brain hemispheres were homogenized and extracted serially in Tris-buffered saline (TBS), TBS plus 1% Triton X-100 (TBS-T), guanidine buffer (5 M guanidine HCl/50 mM Tris, pH 8.0) as described in our previous study (Liu Y et al. 2014). Aβ concentrations in three separate fractions of brain homogenates were determined by Invitrogen™ Amyloid β 42 and 40 Human ELISA kits (Cat.-No.: KHB3441 and KHB3481, respectively; both from Thermo Fisher Scientific). Results were normalized on the basis of the sample’s protein concentration.

### Quantitative PCR for analysis of gene transcripts

Total RNA was isolated from mouse brains with TRIzol or from selected CD11b or CD4-positive cells with RNeasy Plus Mini Kit (Qiagen) and reverse-transcribed. Gene transcripts were quantified with established protocols (Liu Y et al. 2014; Hao et al. 2016) and Taqman gene expression assays of mouse *Tnf-*α, *Il-1*β, *Chemokine (C–C motif) ligand 2* (*Ccl-2*), *Il-10*, *Chitinase-like 3* (*Chi3l3*), *Mannose receptor C type 1* (*Mrc1*), *Apoe*, *Trem2*, *P2ry12*, *Cx3cr1*, *Lpl*, *Clec7a*, *Itgax*, *Il-17a*, *Ifn-*γ, *Il-4*, *neprilysin* and *insulin-degrading enzyme* (*Ide*) and *Gapdh* (Thermo Fisher Scientific).

### Statistical analysis

Data were presented as mean ± *SEM*. For multiple comparisons, we used one-way or two-way ANOVA followed by Bonferroni, Tukey, or Dunnett T3 *post hoc* test (dependent on the result of Levene’s test to determine the equality of variances). Two independent-samples Students *t*-test was used to compare means for two groups of cases. All statistical analyses were performed with GraphPad Prism 8 version 8.0.2. for Windows (GraphPad Software, San Diego, USA). Statistical significance was set at p < 0.05.

## Results

### Deletion of gut bacteria reduces Il-17a-expressing CD4-positive lymphocytes in APP-transgenic mice

To investigate the relationship between gut and brain, we deleted intestinal bacteria with a published protocol (Liu S et al. 2016) by feeding 3-month-old APP-transgenic mice with an antibiotic cocktail (ampicillin, vancomycin, neomycin, streptomycin, and metronidazole) in the drinking water for 2 months. The colon was enlarged (data not shown), as seen by other groups (Erny et al. 2015). The α-diversity analysis of intestinal bacteria showed that Sobs, Ace, Chao, and Shannon indexes decreased and Simpson index increased in antibiotic-treated APP-transgenic mice compared to APP-transgenic littermates with normal drinking water (Supplementary Fig. 1, A - E; *t* test, *p* < 0.05), indicating that the bacterial richness and diversity in the gut of AD mice were significantly reduced by antibiotic treatments. The β-diversity-based PCoA analysis clearly showed the difference in the intestinal bacterial architecture between APP-transgenic mice with and without antibiotic treatments (Supplementary Fig. 1, F; ANOSIM, R = 1.0000, *p* = 0.008). By comparing proportions of bacterial components on phylum level in these two groups, we observed that antibiotic treatment dramatically reduced bacteria of *Bacteroidetes* and *Firmicutes* phyla (Supplementary Fig. 1, G; Wilcoxon rank-sum test, *p* < 0.05). Bacteria of the *Bacteroidetes* phylum produce high levels of acetate and propionate, while bacteria of the *Firmicutes* phylum produce high amounts of butyrate (Macfarlane and Macfarlane 2003). It should be noted that the bacteria of *Proteobacteria* phylum could be resistant and were even enriched after antibiotic treatments (Supplementary Fig. 1, G; Wilcoxon rank-sum test, *p* < 0.05).

Our recent study showed that Il-17a-expressing CD4-positive lymphocytes increase in the gut and spleen of APP-transgenic mice (Luo et al. 2022). We hypothesized that circulating T cells mediate the effects of intestinal bacteria on microglial activation. CD4-positive cells were selected from the spleen of APP-transgenic mice and quantified for gene transcripts. As shown in Fig. 1, A - D, deletion of gut bacteria by antibiotic treatment for 2 months significantly decreased the transcription of *Il-17a*, but increased the transcription of *Ifn-*γ and *Il-4* (*t* test, *p* < 0.05). We further treated APP-transgenic mice expressing Il-17a-eGFP reporter (Esplugues et al. 2011) with and without antibiotics, and observed that deletion of gut bacteria for 2 months significantly reduced eGFP-expressing CD4+ lymphocytes in both lamina propria and Peyer’s patches of the gut compared to control AD mice with normal drinking water (Fig. 1, E - H; *t* test, *p* < 0.05). Therefore, deletion of intestinal bacteria leads to a significant reduction in Il-17a-expressing CD4-positive T lymphocytes in AD mice. In our APP-transgenic animal models, expression of GFP was not detected in γδ T lymphocytes in the gut (Luo et al. 2022).

**Figure 1.**
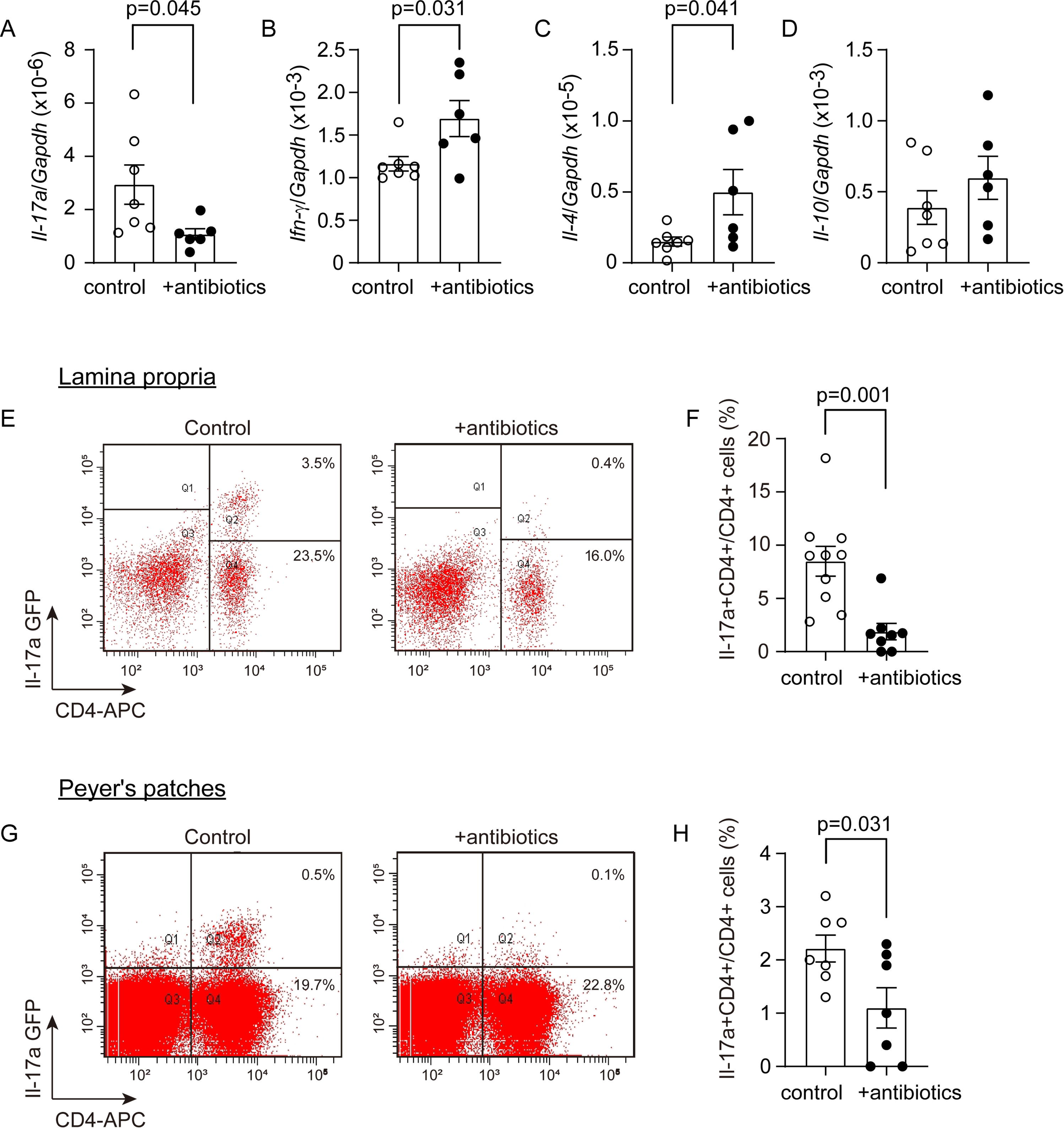
Deletion of gut bacteria reduces Il-17a-expressing CD4-positive lymphocytes. Three-month-old APP-transgenic female mice, expressing or not IL-17a-eGFP reporter, were treated with and without antibiotics in drinking water for 2 months. CD4-positive splenocytes were then selected and detected for transcription of T lymphocyte marker genes. The transcription of *Il-17a* gene was significantly down-regulated by antibiotic treatment, while the transcription of *Ifn-*γ and *Il-4* genes was up-regulated in CD4+ splenocytes compared to APP-transgenic control mice with normal drinking water (A - C; *t* test, *n* ≥ 6 per group). The transcription of *Il-10* gene in spleen cells was not changed by antibiotic treatment (D; *t* test, *n* ≥ 6 per group). In addition, single cell suspensions were prepared from both lamina propria and Peyer’s patches of 5-month-old APP-transgenic mice and analyzed by flow cytometry after fluorescent staining of CD4 (E and G). The expression of IL-17a-associated eGFP was decreased in CD4+ lymphocytes in the intestine of APP-transgenic mice by antibiotic treatments (F and H; *t* test, *n* ≥ 7 per group).

### Deletion of intestinal bacteria inhibits proinflammatory activation in the brain of APP-transgenic mice, but not in Il-17a-deficient AD mice

Deletion of intestinal bacteria has been shown to suppress inflammation in the brain of APP-transgenic mice (Minter et al. 2016; Harach et al. 2017). We did observe that intestinal antibiotic treatment for 2 months significantly reduced the number of Iba1-positive microglia in the hippocampus of APP-transgenic mice from 17.13 ± 1.29 × 10^3^ cells/mm^3^ to 11.85 ± 0.73 × 10^3^ cells/mm^3^ (Fig. 2, A and B; *t* test, *p* = 0.002). Interestingly, in Il-17a-deficient APP-transgenic mice, deletion of gut bacteria did not alter the number of Iba-1-positive cells (Fig. 2, A and C; 16.21 ± 0.57 × 10^3^ cells/mm^3^ vs. 15.85 ± 0.83 × 10^3^ cells/mm^3^; *t* test, *p* > 0.05). We measured transcripts of proinflammatory genes (*Tnf-*α, *Il-1*β, and *Ccl-2*) and antiinflammatory genes (*Il-10*, *Chi3l3*, and *Mrc1*) in brains of APP-transgenic mice. As shown in Fig. 2, E - G, deletion of intestinal bacteria down-regulated the transcription of *Il-1*β and *Ccl-2*, but up-regulated *Il-10* transcription in Il-17a-wildtype APP-transgenic mice (*t* test, *p* < 0.05). Intestinal bacterial deletion did not change the transcription of *Tnf-*α, *Chi3l3* and *Mrc-1* in APP-transgenic mice (Fig. 2, D, H and I; *t* test, *p* > 0.05). In Il-17a-deficient AD mice, deletion of intestinal bacteria did not modulate the transcription of *Tnf-*α, *Il-1*β, *Ccl-2*, *Il-10*, and *Mrc-1* (Fig. 2, D - G, and I; *t* test, *p* > 0.05), except decreasing the transcription of *Chi3l3* (Fig. 2, H; *t* test, *p* < 0.05).

**Figure 2.**
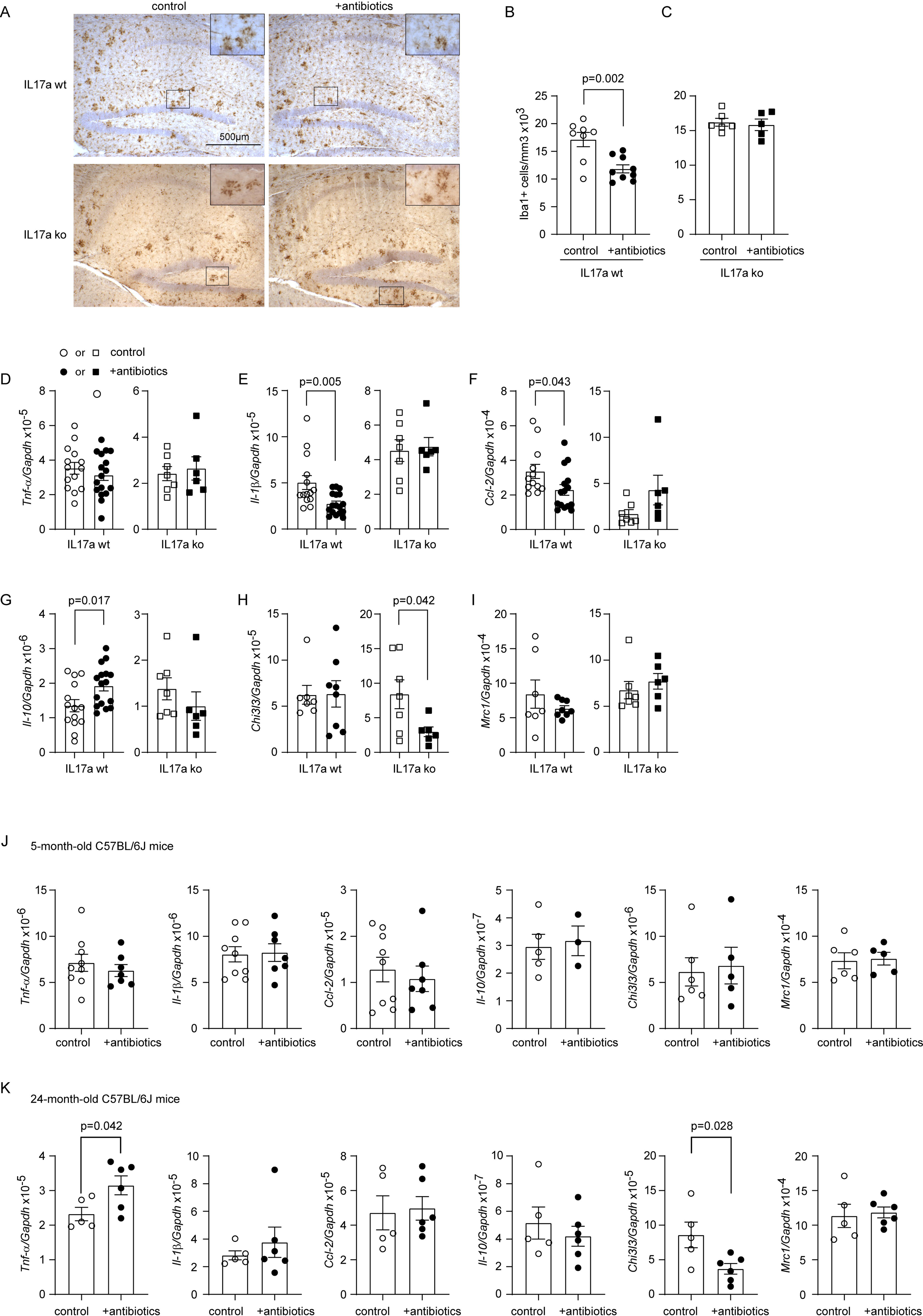
Deletion of gut bacteria reduces inflammatory activation in the brain of Il-17a-wildtype, but not Il17a-deficient APP-transgenic mice. Three-month-old APP-transgenic female mice with (ko) and without (wt) knockout of *Il-17a* gene, and 3 or 22-month-old C57B/L6 female mice were treated with and without antibiotics in drinking water for 2 months. Thereafter, brain tissues were sectioned and microglia were counted with stereological method, Optical Fractionator, after immunohistochemical staining of Iba1 (in brown color) (A - C), or homogenized for RNA isolation and measurement of inflammatory gene transcripts with real-time PCR (D - K). Deletion of gut bacteria significantly decreased Iba1-positive microglia in the hippocampus of Il-17a-wildtype, but not in Il17a-deficient APP-transgenic mice (B and C; *t* test, *n* ≥ 5 per group). Similarly, deletion of gut bacteria significantly reduced the transcripts of *Il-1*β and *Ccl-2*, and increased *Il-10* transcript in the brain of Il-17a-wildtype, but not in Il17a-deficient APP-transgenic mice (E, F, and G; *t* test, *n* ≥ 6 per group). However, deletion of gut bacteria significantly reduced the transcript of *Chi3l3* in the brain of Il17a-deficient APP-transgenic mice (H; *t* test, *n* ≥ 6 per group). Moreover, deletion of gut bacteria did not change the transcription of various inflammatory genes (*Tnf-*α, *Il-1*β, *Ccl-2*, *Il-10*, *Chi3l3*, and *Mrc1*) in in the brain of 5-month-old C57BL/6J female mice (J; *t* test, *n* ≥ 3 per group), however, significantly increased *Tnf-*α transcription and decreased *Chi3l3* transcription in 24-month-old C57B/L6 female mice (K; *t* test, *n* ≥ 5 per group).

In order to investigate whether the intestinal bacterial deletion-induced inflammatory inhibition was specific for mice with AD pathology, we treated 3 and 22-month-old C57BL/6J mice with and without antibiotics for 2 months. Deletion of intestinal bacteria did not change transcription of all genes tested in 5-month-old C57BL/6J mice (*Tnf-*α, *Il-1*β, *Ccl-2*, *Il-10*, *Chi3l3*, and *Mrc1*; Fig. 2, J; *t* test, *p* > 0.05). In 24-month-old C57BL/6J mice, deletion of intestinal bacteria increased *Tnf-*α transcription and decreased *Chi3l3* transcription (Fig. 2, K; *t* test, *p* < 0.05). These experiments also suggested that it was intestinal bacterial deletion instead of antibiotics themselves modifying the inflammatory activation in the brain of APP-transgenic mice.

### Deletion of intestinal bacteria inhibits microglial activation in the brain of APP-transgenic mice, which is abolished by knockout of *Il-17a* gene

To understand the mechanism, through which intestinal antibiotic treatment modified neuroinflammation in AD mice, we selected CD11b^+^ brain cells from 5-month-old APP-transgenic mice with and without 2-month treatments of antibiotics and detected transcripts of disease-associated microglia (DAM) signature genes (Keren-Shaul et al. 2017). Deletion of intestinal bacteria significantly reduced transcription of *Tnf-*α, *Il-1*β, *Ccl-2*, and *Il-10* genes in cells derived from Il-17a-wildtype APP-transgenic mice (Fig. 3, A -D; *t* test, *p* < 0.05), but not in cells from Il-17a-deficient control AD mice. Deletion of intestinal bacteria even increased transcription of *Il-10* gene in Il-17a-deficient APP-transgenic mice (Fig. 3, D; *t* test, *p* < 0.05). Gut bacterial deletion down-regulated transcription of *Apoe* gene in both Il17a-deficient and wildtype APP-transgenic mice (Fig. 3, E; *t* test, *p* < 0.05). Antibiotic treatment did not significantly change the transcription of other genes (*Trem2*, *Cx3cr1*, *P2ry12*, *Clec7a*, *Lpl* and *Itgax*) tested in both Il-17a-deficient and wild-type mice (Fig. 3, F - K; *t* test, *p* > 0.05).

**Figure 3.**
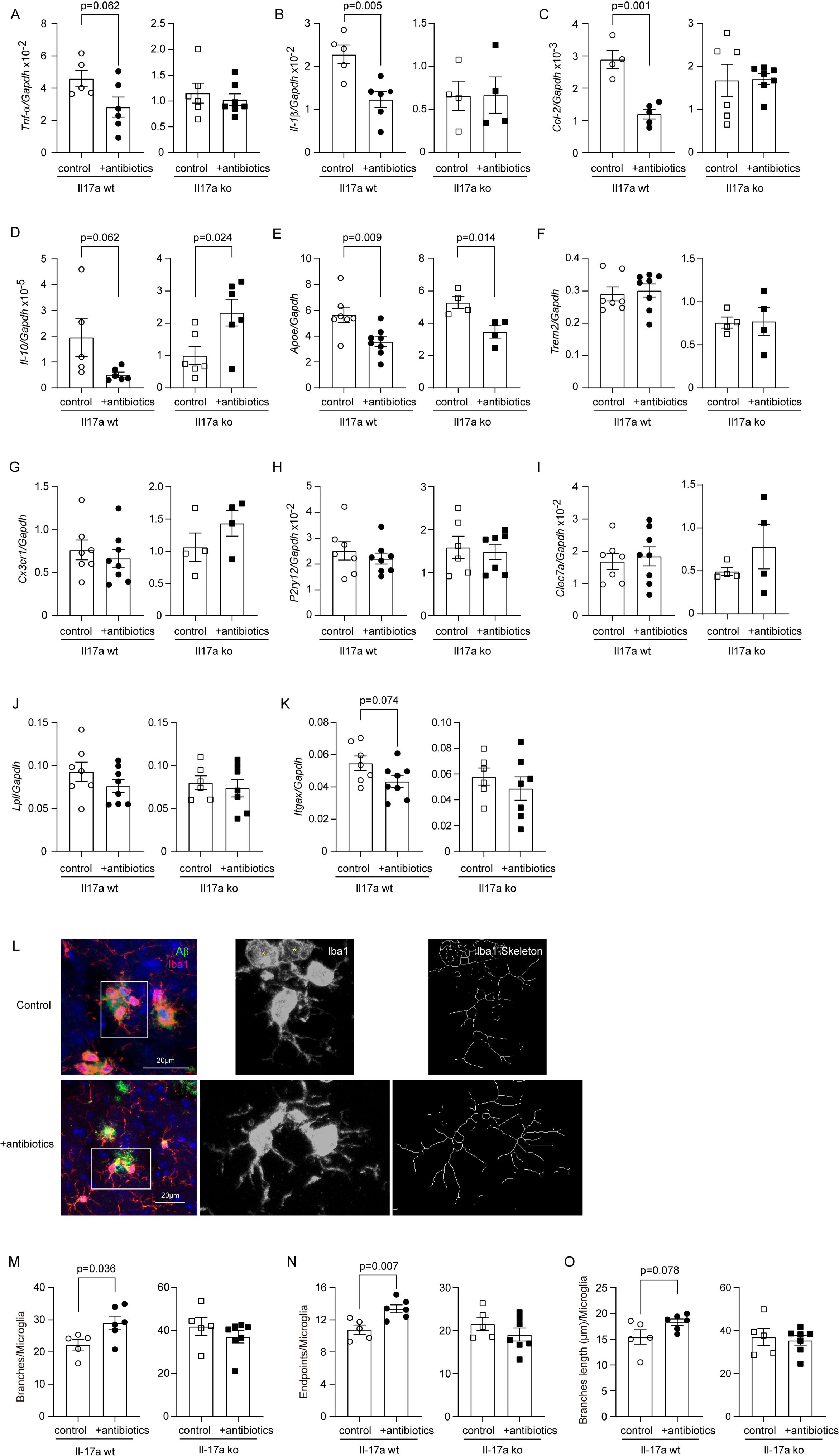
Deletion of gut bacteria inhibits inflammatory activation in microglia in the brain of Il-17a-wildtype, but not Il17a-deficient APP-transgenic mice. Three-month-old APP-transgenic female mice with (ko) and without (wt) knockout of *Il-17a* gene were treated with and without antibiotics in drinking water for 2 months. CD11b-positive brain cells were selected and quantified for the transcription of DAM marker genes. Deletion of gut significantly decreased the transcription of *Tnf-*α, *Il-1*β, *Ccl-2*, and *Il-10* genes, and intended to down-regulate the *Itgax* gene transcription in the brain of Il-17a-wildtype, but not in Il17a-deficient APP-transgenic mice (A - D, and K; *t* test, *n* ≥ 4 per group). Deletion of gut significantly reduced transcription of *Apoe* gene, but not *Trem2*, *Cx3cr1*, *P2ry12*, *Clec7a*, and *Lpl* genes in both Il-17a-wildtype and deficient APP-transgenic mice (E - J; *t* test, *n* ≥ 4 per group). In the analysis of microglial morphology after immunofluorescent staining of Iba1 (L), deletion of gut bacteria increased the number of branches and endpoints of the processes of microglia in Il-17a-wildtype, but not in Il17a-deficient APP-transgenic mice (M - O; *t* test, *n* ≥ 5 per group). The Iba1-positive celles marked with “*” without showing clear processes were excluded for analysis of microglial morphology (L).

In further experiments, we labelled microglia with Iba-1 antibodies and analyzed the morphology of microglia in the vicinity of Aβ deposits as we had done previously (Luo et al. 2022). Deletion of intestinal bacteria increased the total number and end points of branching microglial processes (Fig. 3, L - N; *t* test, *p* < 0.05), and tended to increase branch length (Fig. 3, O; *t* test, *p* = 0.078) in 5-month-old APP-transgenic mice, consistent with previous observations (Erny et al. 2021; Xie et al. 2023). However, all changes in microglial morphology caused by gut bacteria deletion were absent in Il-17a-deficient AD mice (Fig. 3, M - O; *t* test, *p* > 0.05).

### Deletion of intestinal bacteria reduces cerebral A**β** in Il-17a-wildtype but not in Il-17a-deficient APP-transgenic mice

Extracellular Aβ plaques are a pathological hallmark of AD. Deletion of gut bacteria was reported to attenuate Aβ deposits in APP-transgenic mice (Minter et al. 2016; Dodiya et al. 2019). We treated 3-month-old Il-17a-wild-type and deficient female APP-transgenic mice with an antibiotic cocktail for 2 months. As shown in Fig. 4, A - C, 2-month treatments with antibiotics significantly reduced immunoreactive Aβ density from 6.02% ± 0.32% to 4.42% ± 0.64% in the cortex and from 5.67 % ± 0.34 % to 4.16 % ± 0.58% in the hippocampus (*t* test, *p* < 0.05) of Il-17a-wildtype AD mice. In Il-17a-deficient APP-transgenic mice, treatment with antibiotics changed the density of Aβ in neither hippocampus nor cortex (Fig. 4, D - F; *t* test, *p* > 0.05). The brain section was also stained with methoxy-XO4 that typically binds to the β sheet structure of Aβ aggregates. Similarly, treatments with antibiotics significantly reduced Aβ deposits in the cortex (Fig. 4, G and H; *t* test, *p* < 0.05) and tended to decrease Aβ in the hippocampus (Fig. 4, G and I; *t* test, *p* = 0.060) of Il-17a-wildtype APP-transgenic mice, but did not change density of methoxy-XO4-stained Aβ deposits in Il-17a-deficient AD mice (Fig. 4, J - L; *t* test, *p* > 0.05).

**Figure 4.**
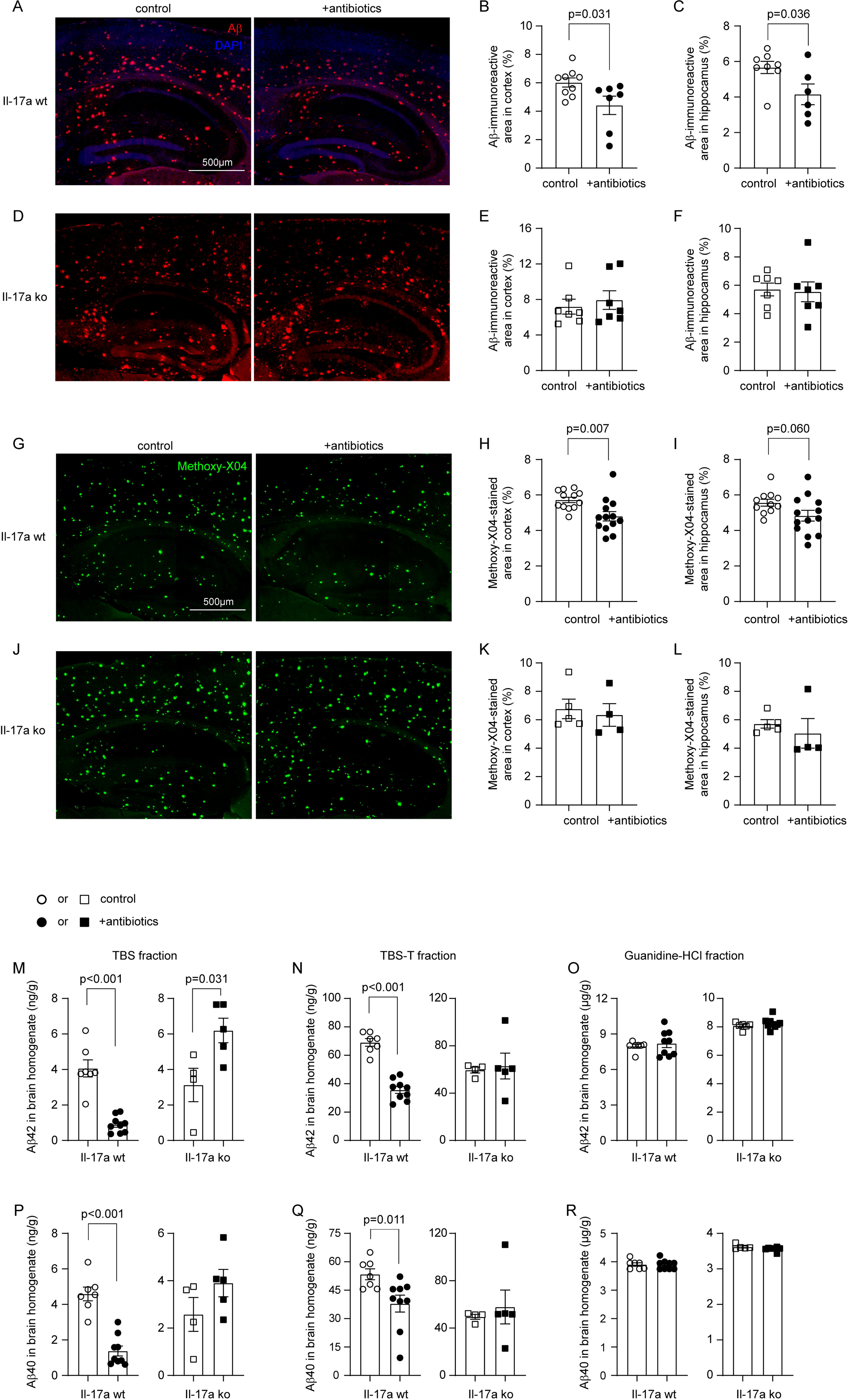
Deletion of gut bacteria reduces Aβ in the brain of Il-17a-wildtype, but not Il17a-deficient APP-transgenic mice. Three-month-old APP-transgenic female mice with (ko) and without (wt) knockout of *Il-17a* gene were treated with and without antibiotics in drinking water for 2 months. Brain tissues were sectioned and stained with immunofluoresce-conjugated Aβ antibodies (A and D) and methoxy-XO4, a fluorescent dye for Aβ aggregates (G and J). Deletion of gut bacteria decreased Aβ deposits in both the hippocampus and cortex after staining of Aβ either with antibody or dye (B, C, H and I; *t* test, *n* ≥ 6 per group). However, in the Il17a-deficient AD mice, deletion of gut bacteria did not change the cerebral Aβ loads (E, F, K and L; *t* test, *n* ≥ 4 per group). Brain tissues were also serially homogenized and extracted in TBS, TBS plus 1% Triton-100 (TBS-T) and guannidine-HCl, and then measured for Aβ40 and Aβ42 with ELISA (M - R). Deletion of gut bacteria decreased the concentrations of both Aβ40 and Aβ42 in TBS- and TBS-T-soluble brain fractions, but not in guanidine-HCl-soluble fraction of Il-17a-wildtype APP-transgenic mice (M - R; *t* test, *n* ≥ 7 per group). Deletion of gut bacteria did not change the concentrations of Aβ40 and Aβ42 in various fractions of bran homogenates of Il-17a-deficient AD mice (N - R; *t* test, *n* ≥ 4 per group), except that it significantly incrased the concentration of Aβ42 in TBS-soluble fraction (M; *t* test, *n* ≥ 4 per group).

We went on measuring Aβ in the brain homogenate with ELISA. In Il-17a wild-type APP-transgenic mice, antibiotic treatment significantly decreased the levels of both Aβ42 and Aβ40 in TBS- and TBS-T fractions (Fig. 4, M, N, P and Q; *t* test, *p* < 0.05), but not in guanidine-soluble fraction (Fig. 4, O and R; *t* test, *p* > 0.05). TBS-, TBS-T-, and guanidine-soluble brain homogenates are enriched with monomeric, oligomeric and high-molecular-weight aggregated Aβ, respectively (Liu Y et al. 2005). In Il-17a-deficient APP-transgenic mice, treatment with antibiotics did not reduce Aβ42 and Aβ40 in all three fractions of brain homogenates (Fig. 4, M - R). On the contrary, antibiotic treatment even increased the concentration of Aβ42 in TBS-soluble fraction of brain homogenate (Fig. 4, M; *t* test, *p* < 0.05).

### Conditional knockout of MyD88 in microglia abolishes intestinal bacterial deletion-induced inflammatory inhibition in the brain of APP-transgenic mice

In this project, we observed that deletion of gut bacteria decreased inflammatory activation and increased the endpoints of microglial branching processes in AD mice, which was abrogated by knockout of *Il-17a* gene. However, we had found that knockout of *Il-17a* gene reduced microglial branching, indicating microglial inflammatory activation, in APP-transgenic mouse brain in a previous study (Luo et al. 2022), which suggested that Il-17a modulated rather than mediated the effect of intestinal bacteria on microglia. We suspected that there were other mechanisms mediating the effect of gut bacteria.

As structural components of bacteria in the gut (e.g., LPS) have the potential to circulate into the brain and directly activate microglia through Toll-like receptors (Dominy et al. 2019; Marizzoni et al. 2023), we hypothesized that MyD88 might regulate microglial activation in AD mice after antibiotic treatment. We continued to treat microglial MyD88-haploinsufficient APP-transgenic mice (APP^tg^MyD88^lox/wt^Cre^+/-^), which we established in a previous study (Quan et al. 2021), with and without antibiotics. In 5-month-old microglial MyD88-deficient APP-transgenic mice, deletion of intestinal bacteria for 2 months neither reduced density of Iba-1-positive cells in the hippocampus (Fig. 5, A and B; 12.41 ± 0.89 × 10^3^ cells/mm^3^ vs. 12.08 ± 0.95 × 10^3^ cells/mm^3^; *t* test, *p* > 0.05), nor changed the transcription of *Tnf-*α, *Il-1*β, *Ccl-2*, *Il-10*, *Chi3l3*, and *Mrc1* in the brain tissue (Fig. 5, C - H; *t* test, *p* > 0.05). Moreover, deletion of intestinal bacteria did not alter the transcription of DAM signature genes (*Tnf-*α, *Il-1*β, *Ccl-2*, *Il-10*, *Apoe*, *Trem2*, *Cx3cr1*, *P2ry12*, *Clec7a*, and *Itgax*) (Fig. 5, I - Q, and S; *t* test, *p* > 0.05) except that it down-regulated the transcription of *Lpl* gene in individual MyD88-deficient microglia (Fig. 5, R; *t* test, *p* < 0.05). Additional experiments also showed that haploinsufficiency of MyD88 abolished the effect of gut bacteria deletion on the number and length of microglial branching processes (Fig. 5, T - W; *t* test, *p* > 0.05).

**Figure 5.**
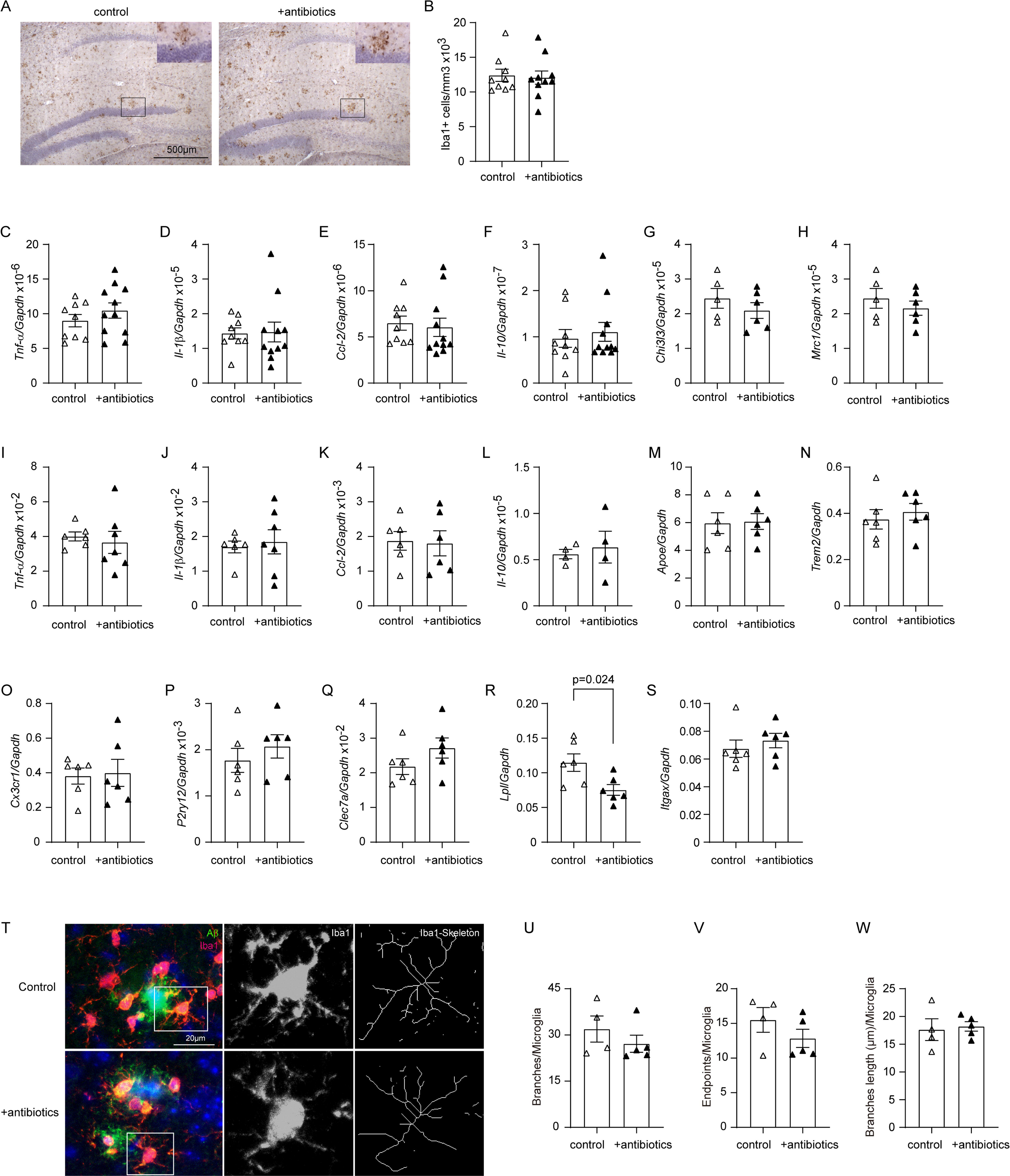
Deletion of gut bacteria does not change the inflammatory activation in the brain of microglial MyD88-haploinsufficient APP-transgenic mice. Three-month-old APP-transgenic female mice with haploinsufficiency of MyD88 in microglia were treated with and without antibiotics in drinking water for 2 months. Neuroinflammation was then assessed either by counting Iba1-positive cells (in brown colour after immunohistochemistry) in the hippocampus (A and B), or by measuring inflammatory gene transcripts in the homogenate of hippocampus and cortex (C - H). CD11b-positive brain cells were also selected and analyzed for the transcriptional levels of DAM genes (I - S). Deletion of gut bacteria did not reduce Iba1-positive microglia, or transcription of inflammatory genes in the brain (A - H; *t* test, *n* ≥ 5 per group). Similarly, deletion of gut bacteria did not alter the transcription of various DAM genes tested, with the exception of decreased transcription of *Lpl* gene (I - S; *t* test, *n* ≥ 4 per group). The morphology of microglia in the vicinity of Aβ deposits was also analyzed after immunostaining with fluorescence-conjugated Iba1 antibodies (T). Deletion of intestinal bacteria did not change the number and length of microlgial processes (U - W; *t* test, *n* ≥ 4 per group).

### Conditional knockout of MyD88 in microglia mitigates intestinal bacterial deletion-induced A**β** reduction in the brain of APP-transgenic mice

After we observed that microglial deficiency of MyD88 abolished the effects of gut bacterial deletion on microglial inflammatory inhibition, we asked whether deficiency of MyD88 in microglia also altered gut bacteria-mediated amyloid modification in AD mouse brain. We stained brain sections of microglial MyD88-haploinsufficient and wild-type APP-transgenic mice with both Aβ antibody and methoxy-X04 fluorescent Aβ dye (Fig. 6, A and D), and observed that antibiotic treatments reduced immunoreactive Aβ deposits in both the cortex and hippocampus (Fig. 6, B and C; *t* test, *p* < 0.05), but did not change the area of methoxy-X04-stained Aβ plaques (Fig. 6, E and F; *t* test, *p* > 0.05). Moreover, deletion of intestinal bacteria decreased the concentration of Aβ42 and Aβ40 in RIPA-soluble brain homogenate in microglial MyD88-wildtype but not MyD88-deficient APP-transgenic mice (Fig. 6, G and H; *t* test, *p* < 0.05).

**Figure 6.**
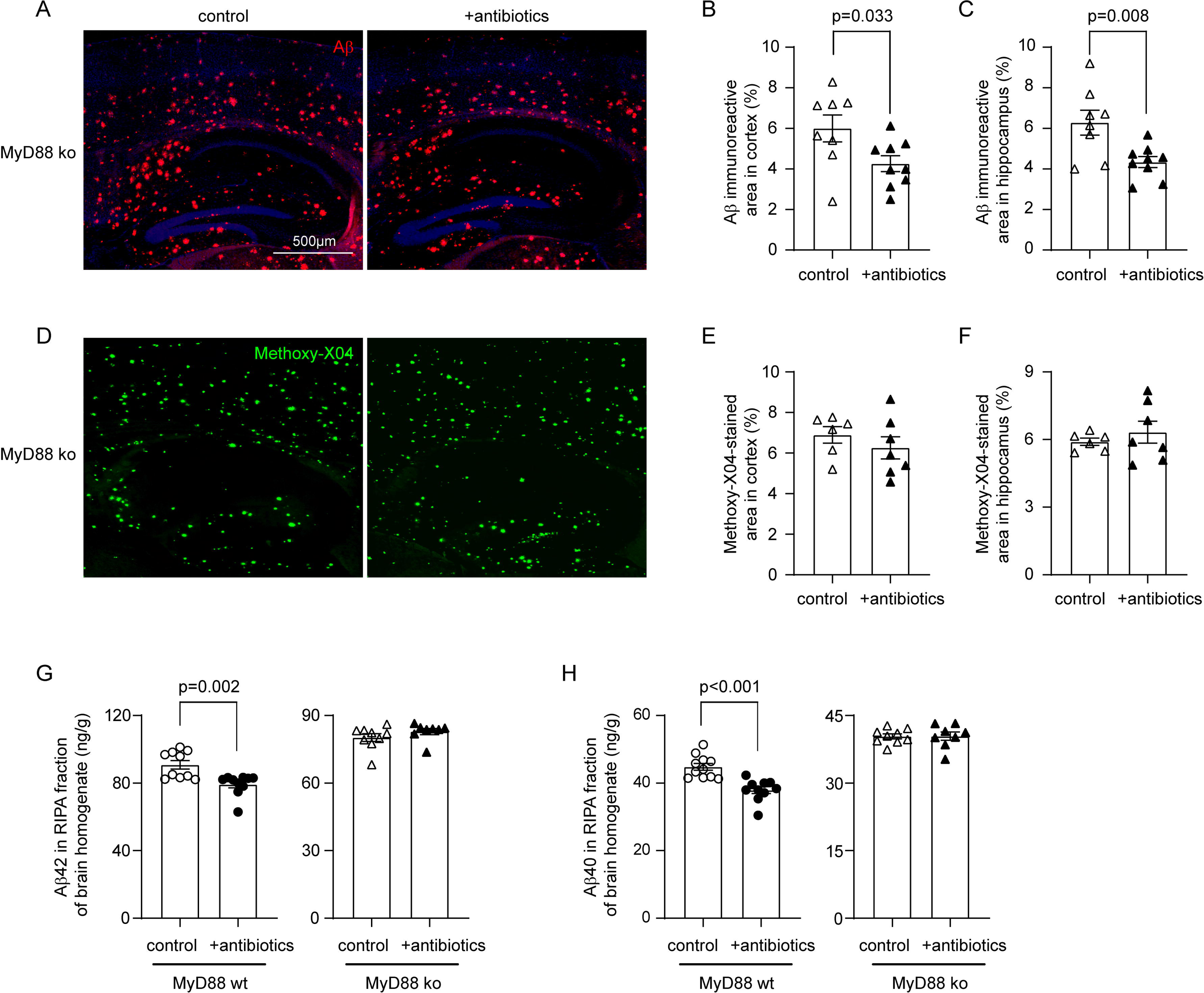
Deletion of gut bacteria reduces Aβ in the brain of Il-17a-wildtype, but not Il17a-deficient APP-transgenic mice. Three-month-old APP-transgenic female mice with (ko) and without (wt) haploinsufficiency of MyD88 in microglia were treated with and without antibiotics in drinking water for 2 months. Brain tissues were stained with immunofluoresce-conjugated Aβ antibodies (A) and methoxy-XO4 (D). Deletion of gut bacteria decreased Aβ deposits in both the hippocampus and cortex after immunofluorescent staining of Aβ, but not after staining with the dye of Aβ aggregates in microglial MyD88-haploinsufficient APP-transgenic mice (B, C, E and F; *t* test, *n* ≥ 6 per group). Brain tissues were also homogenized in RIPA buffer and measured for Aβ40 and Aβ42 with ELISA. Deletion of gut bacteria decreased the concentrations of both Aβ40 and Aβ42 in microglial MyD88-wildtype but not MyD88-haploinsufficient AD mice (G and H; *t* test, *n* ≥ 8 per group).

### Deletion of intestinal bacteria does not increase microglial A**β** phagocytosis and extracellular A**β** degradation, but potentially reduces A**β** production

In following experiments, we asked how deletion of intestinal bacteria attenuated the amyloid pathology in AD mice. Flow cytometric analysis of brain cells after Aβ staining with methoxy-X04 showed that deletion of intestinal bacteria remarkedly decreased both percentage of Aβ-positive CD11b+ brain cells and the mean fluorescence intensity (mFI) of CD11b+ cell population in APP-transgenic mice (Fig. 7, A - C; *t* test, *p* < 0.05), apparently indicating that microglial internalization of Aβ did not contribute to cerebral Aβ clearance.

**Figure 7.**
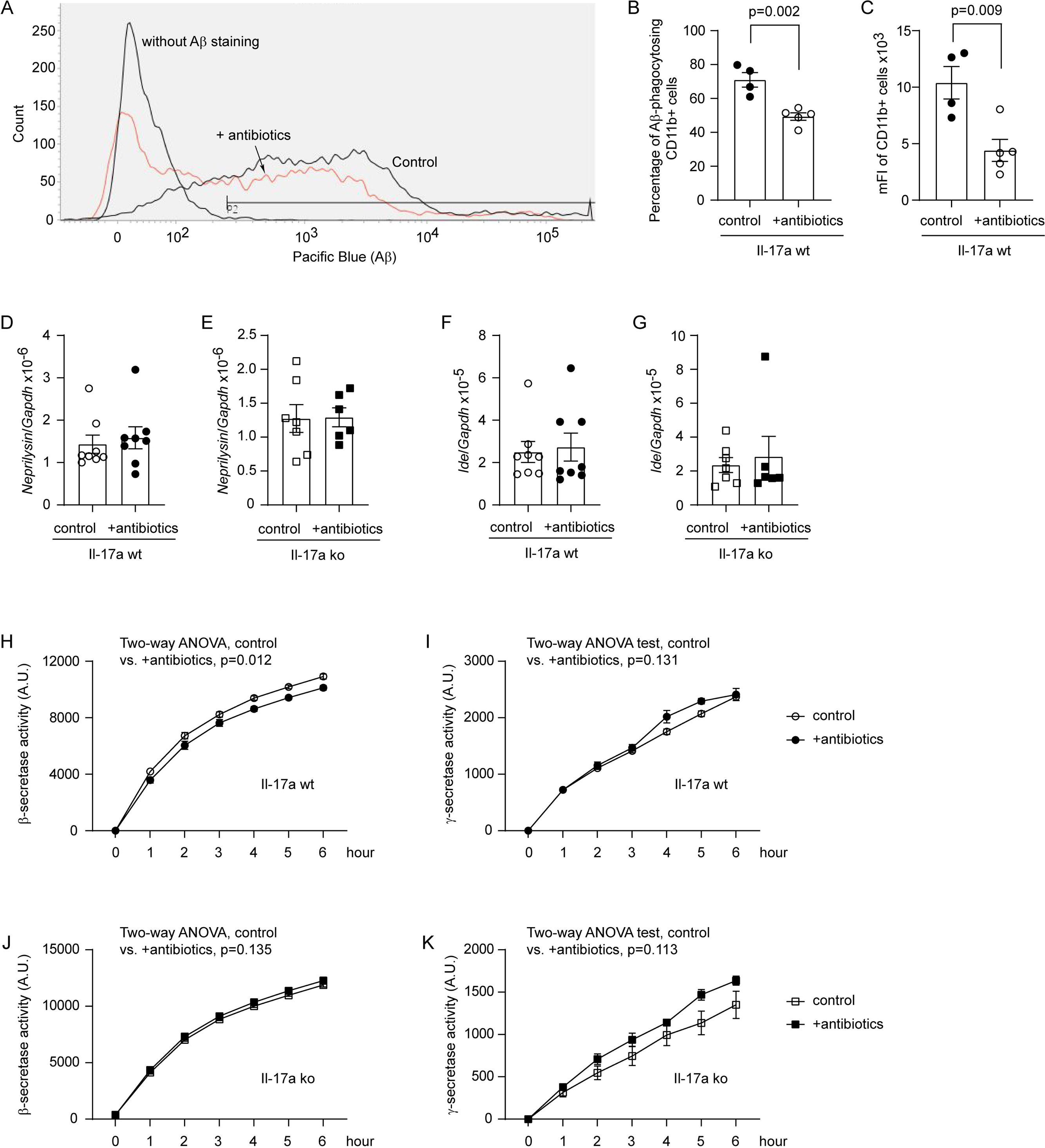
Deletion of gut bacteria reduces β-secretase activity in the brain of Il-17a-wildtype, but not Il17a-deficient APP-transgenic mice. Three-month-old APP-transgenic female mice with (ko) and without (wt) knockout of *Il-17a* gene were treated with and without antibiotics in drinking water for 2 months. After intraperitoneal injection of Aβ dye methoxy-XO4, CD11b-positive brain cells were detected for Aβ-associated fluorescence with flow cytometry (A). Deletion of gut bacteria significantly decreased both the percentage of fluorescent cells among CD11b-positive brain cells, and the mean fluorescence intensity (mFI) of CD11b-positive cell population (B and C; *t* test, *n* ≥ 4 per group). Brain tissues were also collected from AD mice for quantification of *Neprilysin* and *Ide* gene transcripts (D - G), and for β- and γ-secretase assays (H - K). Deletion of gut bacteria did not change the transcription of *Neprilysin* and *Ide* genes (D - G; *t* test, *n* ≥ 4 per group); however, significantly inhibited the activity of β-secretase in the brain of Il-17a-wildtype but not Il17a-deficient APP-transgenic mice (H and J; two-way ANOVA, *n* ≥ 6 per group for Il-17a-wildtype mice, and *n* ≥ 4 per group for Il-17a knockout mice). Deletion of gut bacteria did not change γ-secretase activity in AD mice (I and K; two-way ANOVA, *n* ≥ 6 per group for Il-17a-wildtype mice, and *n* ≥ 4 per group for Il-17a knockout mice).

In the same APP-transgenic mouse strain as we used in this study, another group has reported that the absence of gut bacteria increased protein levels of Aβ-degrading enzymes neprilysin and Ide (Harach et al. 2017). However, our study showed that deletion of intestinal bacteria altered neither *Neprilysin* nor *Ide* gene transcription in the brains of Il-17a-deficient and wildtype APP-transgenic mice (Fig. 7, D - G; *t* test, *p* > 0.05), which suggested that extracellular degradation of Aβ was not the mechanism mediating Aβ reduction in our AD mice.

Our previous study has shown that inhibition of neuroinflammation inhibits β- and γ-secretase activity in the brain of AD mice (Quan et al. 2021). We found that deletion of intestinal bacteria slightly but significantly decreased β-, but not γ-secretase activity (Fig. 7, H and I; two-way ANOVA, antibiotic treatment vs. control, *p* < 0.05), as observed in a published study (Colombo et al. 2021). It was not surprising that the same inhibitory effects of antibiotic treatment on β- and γ-secretase activity could not be seen in Il-17a-deficient APP-transgenic mice (Fig. 7, J and K; two-way ANOVA, antibiotic treatment vs. control, *p* > 0.05), as the antibiotic treatment did not change the inflammatory activation in the brain of Il-17a-deficient AD mice (see Fig. 2).

### Deletion of intestinal bacteria potentially increases A**β** efflux through blood-brain-barrier in APP-transgenic mice, which is driven by Il-17a inhibition

The transportation of Aβ from brain parenchyma to peripheral circulation is an efficient pathway for the cerebral Aβ clearance (Roberts et al. 2014). LRP1 and ABCB1 are a couple of key transporters at the blood-brain-barrier (BBB) that are responsible for Aβ efflux (Kuhnke et al. 2007; Shinohara et al. 2017). We found that the protein levels of Abcb1 and Lrp1were significantly increased in 5-month-old APP-transgenic mice, which had been treated with antibiotics for 2 months (Fig. 8, A, B and D; *t* test, *p* < 0.05). The increase in Abcb1 and Lrp1 induced by gut bacterial deletion was also present in microglial MyD88-haploinsufficient AD mice (Fig. 8, A, C and E; *t* test, *p* < 0.05). Remarkably, the up-regulation of both Lrp1 and Abcb1 in the brain homogenate by deletion of intestinal bacteria was again abolished by knockout of Il-17a gene (Fig. 8, F - H; *t* test, *p* > 0.05).

**Figure 8.**
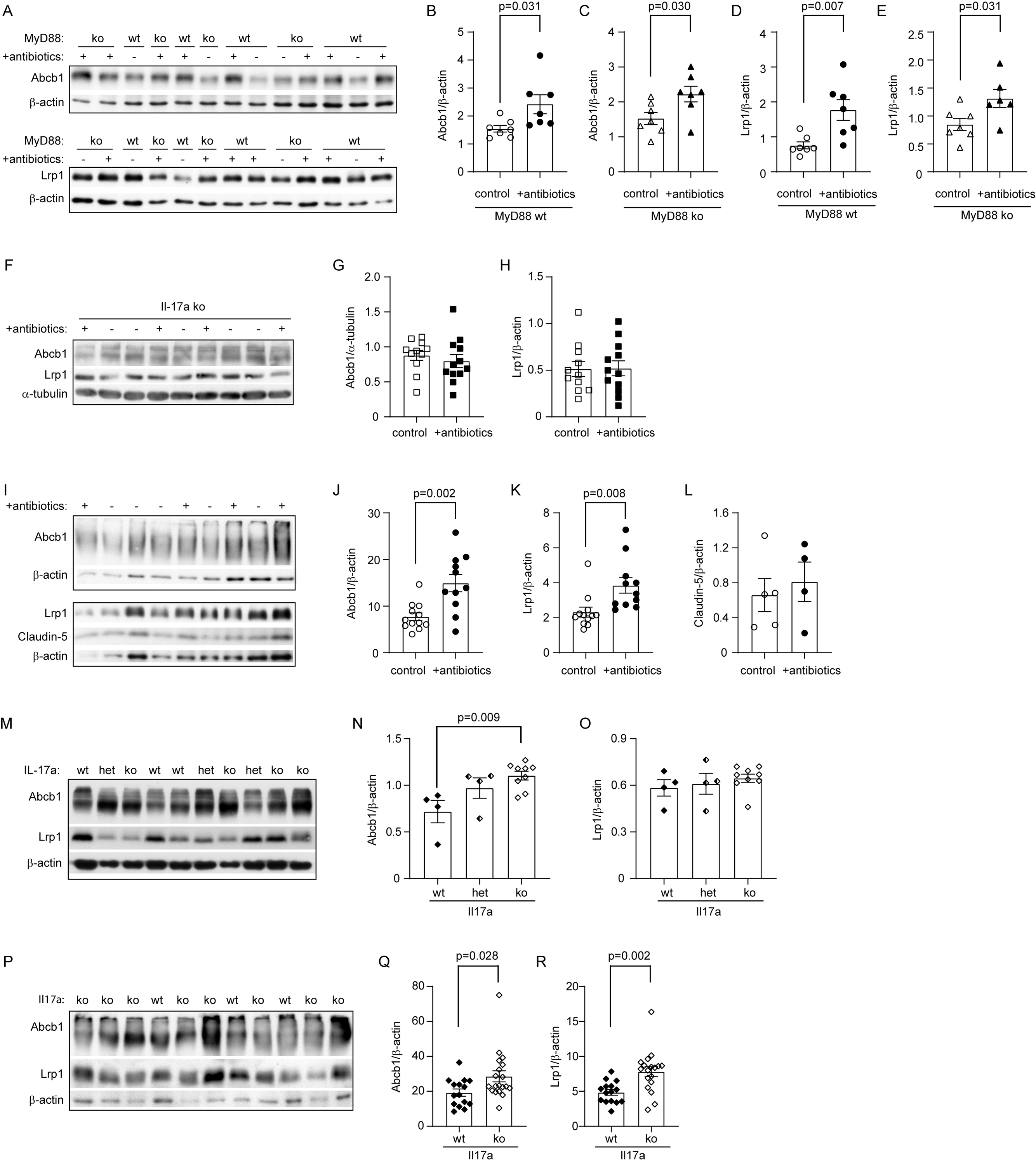
Deletion of gut bacteria increases Abcb1 and Lrp1 expression in the blood-brain-barrier of Il-17a-wildtype or microglial MyD88-deficient, but not Il17a-deficient APP-transgenic mice. Three-month-old APP-transgenic female mice with (ko) and without (wt) haploinsufficiency of MyD88 in microglia were treated with and without antibiotics in drinking water for 2 months. Brain homogenates were quantified for protein levels of Abcb1 and Lrp1 with Western blot (A). Deletion of gut bacteria significantly increased expression of Abcb1 and Lrp1 in brains of both microglial MyD88-wildtype and deficient APP-transgenic mice (B - E; *t* test, *n* = 7 per group for MyD88-wildtype mice, and *n* ≥ 6 per group for MyD88-deficient mice). Cerebral vessels were also isolated from MyD88-wildtype AD mice and detected for Abcb1 and Lrp1 at the BBB. Deletion of gut bacteria also significantly increased the expression of Abcb1 and Lrp1 in blood vessels of APP-transgenic mice (F - H; *t* test, *n* ≥ 11 per group); however, it did not change the protein level of claudin-5 (F and I; *t* test, *n* ≥ 4 per group). Brain homogenates were also prepared from Il-17a-deficient APP-transgenic mice. The protein level of neither Abcb1 nor Lrp1 was changed by deletion of gut bacteria (J - L; *n* ≥ 11 per group). In further experiments, Abcb1 and Lrp1 were detected with quantitative Western blot in the brain homogenates from 5-month-old non-APP-transgenic female mice with different expression of Il-17a (M - O; wild-type [wt], heterozygote [het] and knockout [ko]), and in the isolated blood vessels from 5-month-old Il17a-wt and ko APP-transgenic female mice (P - R). Deficiency of Il-17a significantly inceased Abcb1 but not Lrp1 in the non-APP-transgenic mouse brain in a gene dose-dependent manner (N; one-way ANOVA followed by Bonferroni *post-hoc* test, *n* ≥ 4 per group), and increased protein levels of both Abcb1 and Lrp1 in the blood vessels of APP-transgenic mice (Q and R; *t* test, *n* ≥ 15 per group).

We also isolated blood vessels from 5-month-old APP-transgenic mice with and without treatments with antibiotics. We validated the results from the entire brain homogenate that treatments with antibiotics strongly elevated protein levels of Abcb1 and Lrp1 at the BBB (Fig. 8, I - K; *t* test, *p* < 0.05). Notably, the protein level of claudin-5 in cerebral blood vessels did not differ between APP-transgenic mice with and without antibiotic treatment (Fig. 8, I and L; *t* test, *p* > 0.05). Similarly, we did not detect any mouse IgG in brain homogenates of antibiotics-treated AD mice with Western blot (data not shown), which suggests that deletion of intestinal bacteria increased expression of Abcb1 and Lrp1 at the BBB, but did not significantly impair the BBB.

In further experiments, we detected Lrp1 and Abcb1 in the brain homogenates from 5- month-old APP-non-transgenic female mice with different expression of Il-17a. We observed that deficiency of Il-17a significantly increased the protein level of Abcb1, but not Lrp1, in a gene dose-dependent way (Fig. 8, M - O; one-way ANOVA, *p* < 0.05). Similarly, we isolated blood vessels from Il-17a-deficient and wild-type APP-transgenic mice and observed that deficiency of Il-17a significantly increased both Lrp1 and Abcb1 in the tissue lysate of blood vessels (Fig. 8, P - R; *t* test, *p* < 0.05), suggesting that deletion of intestinal bacteria potentially increases Lrp1/Abcb1-mediated Aβ efflux through inhibiting Il-17a signaling.

## Discussion

Gut bacteria contribute to AD development (Chandra et al. 2023), but how gut bacteria regulate brain pathology remains unclear. By using young female APP-transgenic mice with rapidly developing Aβ pathology, we found that deletion of gut bacteria decreased microglial inflammatory activation and brain Aβ levels, particularly soluble Aβ. This effect was reversed by knockout of *Il-17a* gene or haploinsufficiency of MyD88 specifically in microglia. The attenuation of cerebral Aβ pathology by gut bacterial deletion may be attributed to the inhibition of β-secretase activity and the upregulation of Aβ efflux-related Lrp1 and Abcb1 expression in the brain, which was also abrogated by Il-17a deficiency.

Bacterial phenotypes are potentially transferred between contacting mice (Zhang Y et al. 2023). We housed 4-6 female mice in a cage for treatments with and without antibiotics, which might reduce the variability of gut microbiota-induced brain responses between mice within the same experimental group. We observed that deletion of gut bacteria in 5-month-old APP-transgenic mice inhibited proinflammatory activation in the brain as evidenced by reduction of microglial number, down-regulation of *Il-1*β and *Ccl-2* transcription and up- regulation of *Il-10* transcription, which are consistent with published studies (Harach et al. 2017; Minter et al. 2017; Colombo et al. 2021). However, we noted that intestinal bacterial deletion in 5-month-old C57BL/6J mice did not have the same effect on the inflammatory modulation as in AD mice, and even promoted inflammatory activation (e.g., transcriptional up-regulation of *Tnf-*α, and down-regulation of *Chi3l3*) in the brain of 24-month-old C57BL/6J mice. Since transplantation of intestinal bacteria from old mice to young mice increases the permeability of the colonic mucosa, indicating a change in the structure and function of the gastrointestinal tract during aging (Zeng et al. 2023), it stands to reason that a significant inflammatory modulation caused by gut bacterial deletion depends on the priming status and pattern of inflammatory brain cells (e.g., microglia).

We observed that deletion of gut bacteria inhibited the development of Il-17-expressing CD4-positive cells, while Th1 and Th2 cells potentially increased, which aligns with previous studies (Lee et al. 2011; Benakis et al. 2016; Minter et al. 2017). Given that IL-17a promotes tissue inflammation (Xu and Cao 2010), we hypothesized that Il-17a mediates the impact of intestinal bacterial deletion on brain cells. Indeed, the deficiency of Il-17a abolished the inflammatory inhibition induced by the deletion of gut bacteria in both microglia and brain tissue. However, knockout of *Il-17a* gene reduces microglial branching in APP-transgenic mice, indicating inflammatory activation of microglia, as shown in our recent project (Luo et al. 2022). Thus, Il-17a regulates rather than mediates the effects of gut bacteria on the brain.

In AD patients, bacterial components such as LPS and gingipains increase in the blood and are co-localized with Aβ plaques (Zhan et al. 2016; Zhao et al. 2017; Dominy et al. 2019; Marizzoni et al. 2023). In APP-transgenic mice, oral infection with *P. gingivalis* leads to infection of the brain and induction of Aβ42 (Dominy et al. 2019). We thus hypothesized that microglia could be directly primed by bacterial structure components. Our experiments showed that haploinsufficiency of MyD88 abolished the inflammatory inhibition in microglia caused by the deletion of gut bacteria in APP-transgenic mice. A lack of MyD88 might minimize the inflammatory priming of microglia. The deletion of gut bacteria cannot exert an additional inhibitory effect on microglia. Alternatively, the deletion of gut bacteria could produce an inhibitory molecule that actively suppresses the inflammatory activation of microglia through MyD88, but this would require further proof. It should be noted that metabolites of gut bacteria, such as SCFAs, do not always have anti-inflammatory properties (Maslowski et al. 2009). Some of them, e.g., valeric acid, have been shown to increase the concentrations of IL-17, IL-1β, and IL-6 in the blood and brain of mice with experimental stroke (Zeng et al. 2023). A study in elderly people with cognitive performance ranging from normal to dementia showed that valeric acid and acetic acid together with LPS and proinflammatory cytokines in the blood were correlated with the severity of amyloid pathology in the brain (Marizzoni et al. 2020). It remains to be investigated whether SCFAs signaling is also regulated by MyD88.

Previous research (Dodiya et al. 2019) has shown that oral administration of antibiotics at a high dose for 14 - 21 days after birth, followed by a low dose thereafter, reduced Aβ deposition in the brains of male but not female APP-transgenic mice. Perturbation of gut bacteria had no effect on soluble Aβ. However, our study found that treating female APP-transgenic mice with antibiotics at a high dose from 3 to 5 months of age resulted in a decrease in Aβ load, especially soluble Aβ, in the brain. Furthermore, both studies showed a similar favorable effect of antibiotic treatment on the homeostatic status of microglia in APP-transgenic mice. Despite the differences in antibiotic treatment protocols, we do not consider sex to be the primary factor influencing the effects of gut bacteria on Aβ pathology. Moreover, deletion of gut bacteria, particularly in *Bacteroidetes* and *Firmicutes* phyla, dramatically reduced Aβ phagocytosis in our APP-transgenic mice, confirming the essential role of gut bacteria-producing SCFAs in the maturation and activation of microglia (Erny et al. 2015; Colombo et al. 2021; Erny et al. 2021). Our experiment indicated that microglia are not responsible for the cerebral Aβ reduction in APP-transgenic mice after deletion of gut bacteria.

Transportation of Aβ from brain parenchyma to peripheral plasma might contribute 25% of total clearance of cerebral Aβ (Roberts et al. 2014). LRP1 and ABCB1 are two key transporters on blood-brain-barrier (BBB) responsible for Aβ efflux (Kuhnke et al. 2007; Shinohara et al. 2017). Deletion or inhibition of endothelial Lrp1 and Abcb1 results in Aβ accumulation in AD mouse brain (Cirrito et al. 2005; Storck et al. 2016). We observed that deletion of gut bacteria increased the expression of Lrp1 and Abcb1 in APP-transgenic mice. Knockout of *Il-17a* gene increased the expression of Lrp1 and Abcb1 in the BBB of APP-transgenic mice without treatment with antibiotics. Thus, it is plausible that inhibition of Il-17a signaling induces expression of Lrp1 and Abcb1 at BBB, which in turn increases Aβ efflux.

The upregulation of Lrp1 and Abcb1 expression by intestinal bacterial deletion was reversed by knockout of *Il-17a* gene, but not by conditional knockout of *Myd88* in microglia, suggesting that the increase of Lrp1 and Abcb1 is not due to the neuroinflammatory inhibition in AD mice. There is one remaining question as to why Lrp1 and Abcb1 increase in both microglial MyD88-deficient and wild-type APP-transgenic mice after gut bacterial deletion, but the lack of MyD88 in microglia could still abrogate the effects of bacterial deletion on cerebral Aβ reduction. We hypothesized that reduced ApoE expression in microglia due to gut bacterial deletion, which occurred in MyD88 wild-type but not deficient APP-transgenic mice, may be an underlying mechanism. It has been reported that the absence of ApoE retains less Aβ in the brain parenchyma and increases soluble Aβ in the interstitial fluid, possibly promoting the diffusion of soluble Aβ from the parenchyma into the perivascular space (DeMattos et al. 2004), which favors Aβ efflux. Notably, deletion of gut bacteria decreased *Apoe* gene transcription but did not change the cerebral Aβ level in Il-17a-deficient AD mice, which might be due to the failure in up-regulating Lrp1 and Abcb1 at the BBB. Our study indicates the importance of cooperation between microglia and BBB in the clearance of cerebral Aβ.

Aβ is produced by β-(BACE1) and γ-secretases after serial digestions of APP. The published studies by our group and others have shown that the activity or protein level of β- and γ-secretases is regulated by inflammatory activation in the brain (He et al. 2007; Hur et al. 2020; Quan et al. 2021). Not surprisingly, deletion of gut bacteria inhibited β-secretase activity in correlation with inflammatory inhibition in our APP-transgenic mice, which is consistent with the observation in germ-free AD mice (Colombo et al. 2021). We did not detect an increase in the expression of neprilysin and Ide in our APP-transgenic mice after antibiotic treatment, suggesting that extracellular degradation of Aβ is not the key mechanism for Aβ clearance after gut bacterial deletion. This result differs from the observation in a study on germ-free AD mice (Harach et al. 2017).

In summary, our results are consistent with published observations that deletion of gut bacteria attenuates inflammatory activation of microglia and Aβ pathology in the brain of APP-transgenic mice; however, we have further shown that these inhibitory effects of gut bacterial deletion are regulated by the expression of Il-17a and microglial MyD88. One possible mechanism mediating Aβ reduction is that deletion of gut bacteria inhibits Il-17a signaling, e.g., blocks the development of IL-17a-expressing CD4-positive lymphocytes, which increases Lrp1 and Abcb1 expression in BBB and promotes Aβ efflux from the brain parenchyma to circulation. Our study contributes to a better understanding of mechanisms that mediate the gut-brain axis in AD pathogenesis.

## Supporting information

Supplementary Figure 1, Antibiotic treatment dramatically changes the bacterial composition in the gut of APP-transgenic mice.

## Author contributions

Y.L. conceptualized and designed the study, acquired funding, conducted experiments, acquired and analyzed data, and wrote the manuscript. W.H., Q.L., I.T., and W.Q. conducted experiments, acquired data and analyzed data. T.H. provided resource and discussed study design. M.M. provided an animal facility and supervised animal experiments. K.F. provided a research laboratory and supervised the laboratory work. All authors contributed to the article and approved the submitted version.

## Acknowledgments

We thank Prof. M. Jucker (Hertie Institute for Clinical Brain Research, Tübingen) for providing APP/PS1-transgenic mice, Prof. M. Prinz (University of Freiburg) for Cx3Cr1-CreERT2 mice, Prof. Y. Iwakura (Tokyo University of Science) for Il-17a knockout mice and Prof. R. Flavell (Yale University) for Il-17a-eGFP reporter mice. We appreciate Elisabeth Gluding and Kati Jordan for their excellent technical assistance.

## Data availability statement

Our study has not generated new datasets. All data generated or analyzed during this study are included in this published article and its supplementary information files.

---

**Supplementary Figure 1, Antibiotic treatment dramatically changes the bacterial composition in the gut of APP-transgenic mice.**

Three-month-old APP-transgenic female mice were treated with (*n* = 4) and without (*n* = 5) antibiotics in drinking water for 2 months. Intestinal bacteria were harvested from the appendix and colon and sequenced for the V3 -V4 region of 16S rRNA-encoding gene. Using Sobs, Shannon, Ace, Chao and Simpson’s indices, α-diversity analysis shows that treatment with an antibiotic cocktail significantly reduces bacterial richness and diversity within each mouse (A - E; *t* test). Principal coordinate analysis (PCoA) was used for β-diversity analysis of bacterial composition at the genus level in APP-transgenic mice with and without antibiotic treatment (F; Each symbol represents the gut bacteria of an individual mouse). As expected, the structure of gut bacterial community of antibiotics-treated APP-transgenic mice differed significantly from those of APP-transgenic littermates with normal drinking water (F; ANOSIM analysis, +antibiotics vs. control, *p* < 0.05). Bar plots depict abundance (% of total) of the indicated phyla. Wilcoxon rank-sum tests show that antibiotic treatment decreases the relative abudance of bacteria in the phyla *Bacteroidetes* and *Firmicutes*, but increases bacterial abudance in the phyla *Proteobacteria* (G).

## References

Baruch K, Rosenzweig N, Kertser A, Deczkowska A, Sharif AM, Spinrad A, Tsitsou-Kampeli A, Sarel A, Cahalon L, Schwartz M. 2015. Breaking immune tolerance by targeting Foxp3(+) regulatory T cells mitigates Alzheimer’s disease pathology. Nat Commun. 6:7967.

Benakis C, Brea D, Caballero S, Faraco G, Moore J, Murphy M, Sita G, Racchumi G, Ling L, Pamer EG et al. 2016. Commensal microbiota affects ischemic stroke outcome by regulating intestinal gammadelta T cells. Nat Med. 22(5):516–523.

Chandra S, Sisodia SS, Vassar RJ. 2023. The gut microbiome in Alzheimer’s disease: what we know and what remains to be explored. Mol Neurodegener. 18(1):9.

Cirrito JR, Deane R, Fagan AM, Spinner ML, Parsadanian M, Finn MB, Jiang H, Prior JL, Sagare A, Bales KR et al. 2005. P-glycoprotein deficiency at the blood-brain barrier increases amyloid-beta deposition in an Alzheimer disease mouse model. J Clin Invest. 115(11):3285–3290.

Colombo AV, Sadler RK, Llovera G, Singh V, Roth S, Heindl S, Sebastian Monasor L, Verhoeven A, Peters F, Parhizkar S et al. 2021. Microbiota-derived short chain fatty acids modulate microglia and promote Abeta plaque deposition. Elife. 10:e59826.

Couter CJ, Surana NK. 2016. Isolation and Flow Cytometric Characterization of Murine Small Intestinal Lymphocytes. JoVE.(111):e54114.

Dansokho C, Ait Ahmed D, Aid S, Toly-Ndour C, Chaigneau T, Calle V, Cagnard N, Holzenberger M, Piaggio E, Aucouturier P et al. 2016. Regulatory T cells delay disease progression in Alzheimer-like pathology. Brain. 139(Pt 4):1237–1251.

DeMattos RB, Cirrito JR, Parsadanian M, May PC, O’Dell MA, Taylor JW, Harmony JA, Aronow BJ, Bales KR, Paul SM et al. 2004. ApoE and clusterin cooperatively suppress Abeta levels and deposition: evidence that ApoE regulates extracellular Abeta metabolism in vivo. Neuron. 41(2):193–202.

Dodiya HB, Kuntz T, Shaik SM, Baufeld C, Leibowitz J, Zhang X, Gottel N, Zhang X, Butovsky O, Gilbert JA et al. 2019. Sex-specific effects of microbiome perturbations on cerebral Abeta amyloidosis and microglia phenotypes. J Exp Med. 216(7):1542–1560.

Dominy SS, Lynch C, Ermini F, Benedyk M, Marczyk A, Konradi A, Nguyen M, Haditsch U, Raha D, Griffin C et al. 2019. Porphyromonas gingivalis in Alzheimer’s disease brains: Evidence for disease causation and treatment with small-molecule inhibitors. Sci Adv. 5(1):eaau3333.

Dunham SJB, McNair KA, Adams ED, Avelar-Barragan J, Forner S, Mapstone M, Whiteson KL. 2022. Longitudinal Analysis of the Microbiome and Metabolome in the 5xfAD Mouse Model of Alzheimer’s Disease. mBio. 13(6):e0179422.

Erny D, Dokalis N, Mezo C, Castoldi A, Mossad O, Staszewski O, Frosch M, Villa M, Fuchs V, Mayer A et al. 2021. Microbiota-derived acetate enables the metabolic fitness of the brain innate immune system during health and disease. Cell Metab. 33(11):2260–2276 e2267.

Erny D, Hrabe de Angelis AL, Jaitin D, Wieghofer P, Staszewski O, David E, Keren-Shaul H, Mahlakoiv T, Jakobshagen K, Buch T et al. 2015. Host microbiota constantly control maturation and function of microglia in the CNS. Nat Neurosci. 18(7):965–977.

Esplugues E, Huber S, Gagliani N, Hauser AE, Town T, Wan YY, O’Connor W, Jr., Rongvaux A, Van Rooijen N, Haberman AM, et al. 2011. Control of TH17 cells occurs in the small intestine. Nature. 475(7357):514–518.

Goldmann T, Wieghofer P, Muller PF, Wolf Y, Varol D, Yona S, Brendecke SM, Kierdorf K, Staszewski O, Datta M et al. 2013. A new type of microglia gene targeting shows TAK1 to be pivotal in CNS autoimmune inflammation. Nat Neurosci. 16(11):1618–1626.

Grabrucker S, Marizzoni M, Silajdzic E, Lopizzo N, Mombelli E, Nicolas S, Dohm-Hansen S, Scassellati C, Moretti DV, Rosa M et al. 2023. Microbiota from Alzheimer’s patients induce deficits in cognition and hippocampal neurogenesis. Brain. 146(12):4916–4934.

Hao W, Decker Y, Schnoder L, Schottek A, Li D, Menger MD, Fassbender K, Liu Y. 2016. Deficiency of IkappaB Kinase beta in Myeloid Cells Reduces Severity of Experimental Autoimmune Encephalomyelitis. Am J Pathol. 186(5):1245–1257.

Hao W, Liu Y, Liu S, Walter S, Grimm MO, Kiliaan AJ, Penke B, Hartmann T, Rube CE, Menger MD et al. 2011. Myeloid differentiation factor 88-deficient bone marrow cells improve Alzheimer’s disease-related symptoms and pathology. Brain. 134(Pt 1):278–292.

Harach T, Marungruang N, Duthilleul N, Cheatham V, Mc Coy KD, Frisoni G, Neher JJ, Fak F, Jucker M, Lasser T et al. 2017. Reduction of Abeta amyloid pathology in APPPS1 transgenic mice in the absence of gut microbiota. Sci Rep. 7:41802.

He P, Zhong Z, Lindholm K, Berning L, Lee W, Lemere C, Staufenbiel M, Li R, Shen Y. 2007. Deletion of tumor necrosis factor death receptor inhibits amyloid beta generation and prevents learning and memory deficits in Alzheimer’s mice. J Cell Biol. 178(5):829–841.

Hou B, Reizis B, DeFranco AL. 2008. Toll-like receptors activate innate and adaptive immunity by using dendritic cell-intrinsic and -extrinsic mechanisms. Immunity. 29(2):272–282.

Hur J-Y, Frost GR, Wu X, Crump C, Pan SJ, Wong E, Barros M, Li T, Nie P, Zhai Y et al. 2020. The innate immunity protein IFITM3 modulates γ-secretase in Alzheimer’s disease. Nature. 586(7831):735–740.

Kaur H, Nookala S, Singh S, Mukundan S, Nagamoto-Combs K, Combs CK. 2021. Sex-Dependent Effects of Intestinal Microbiome Manipulation in a Mouse Model of Alzheimer’s Disease. Cells. 10(9):2370.

Keren-Shaul H, Spinrad A, Weiner A, Matcovitch-Natan O, Dvir-Szternfeld R, Ulland TK, David E, Baruch K, Lara-Astaiso D, Toth B et al. 2017. A Unique Microglia Type Associated with Restricting Development of Alzheimer’s Disease. Cell. 169(7):1276–1290 e1217.

Kuhnke D, Jedlitschky G, Grube M, Krohn M, Jucker M, Mosyagin I, Cascorbi I, Walker LC, Kroemer HK, Warzok RW et al. 2007. MDR1-P-Glycoprotein (ABCB1) Mediates Transport of Alzheimer’s amyloid-beta peptides--implications for the mechanisms of Abeta clearance at the blood-brain barrier. Brain Pathol. 17(4):347–353.

Lau SF, Wu W, Seo H, Fu AKY, Ip NY. 2021. Quantitative in vivo assessment of amyloid-beta phagocytic capacity in an Alzheimer’s disease mouse model. STAR Protoc. 2(1):100265.

Lee YK, Menezes JS, Umesaki Y, Mazmanian SK. 2011. Proinflammatory T-cell responses to gut microbiota promote experimental autoimmune encephalomyelitis. Proc Natl Acad Sci U S A. 108 Suppl 1:4615–4622.

Liu S, da Cunha AP, Rezende RM, Cialic R, Wei Z, Bry L, Comstock LE, Gandhi R, Weiner HL. 2016. The Host Shapes the Gut Microbiota via Fecal MicroRNA. Cell Host Microbe. 19(1):32-43.

Liu Y, Liu X, Hao W, Decker Y, Schomburg R, Fulop L, Pasparakis M, Menger MD, Fassbender K. 2014. IKKbeta deficiency in myeloid cells ameliorates Alzheimer’s disease-related symptoms and pathology. J Neurosci. 34(39):12982–12999.

Liu Y, Walter S, Stagi M, Cherny D, Letiembre M, Schulz-Schaeffer W, Heine H, Penke B, Neumann H, Fassbender K. 2005. LPS receptor (CD14): a receptor for phagocytosis of Alzheimer’s amyloid peptide. Brain. 128(Pt 8):1778–1789.

Luo Q, Schnoder L, Hao W, Litzenburger K, Decker Y, Tomic I, Menger MD, Liu Y, Fassbender K. 2022. p38alpha-MAPK-deficient myeloid cells ameliorate symptoms and pathology of APP-transgenic Alzheimer’s disease mice. Aging Cell. 21(8):e13679.

Ma C, Li Y, Mei Z, Yuan C, Kang JH, Grodstein F, Ascherio A, Willett WC, Chan AT, Huttenhower C et al. 2023. Association Between Bowel Movement Pattern and Cognitive Function: Prospective Cohort Study and a Metagenomic Analysis of the Gut Microbiome. Neurology. 101(20):e2014–e2025.

Macfarlane S, Macfarlane GT. 2003. Regulation of short-chain fatty acid production. Proc Nutr Soc. 62(1):67–72.

Marizzoni M, Cattaneo A, Mirabelli P, Festari C, Lopizzo N, Nicolosi V, Mombelli E, Mazzelli M, Luongo D, Naviglio D et al. 2020. Short-Chain Fatty Acids and Lipopolysaccharide as Mediators Between Gut Dysbiosis and Amyloid Pathology in Alzheimer’s Disease. J Alzheimers Dis. 78(2):683–697.

Marizzoni M, Mirabelli P, Mombelli E, Coppola L, Festari C, Lopizzo N, Luongo D, Mazzelli M, Naviglio D, Blouin JL et al. 2023. A peripheral signature of Alzheimer’s disease featuring microbiota-gut-brain axis markers. Alzheimers Res Ther. 15(1):101.

Maslowski KM, Vieira AT, Ng A, Kranich J, Sierro F, Di Y, Schilter HC, Rolph MS, Mackay F, Artis D, et al. 2009. Regulation of inflammatory responses by gut microbiota and chemoattractant receptor GPR43. Nature. 461(7268):1282–1286.

Meyer K, Lulla A, Debroy K, Shikany JM, Yaffe K, Meirelles O, Launer LJ. 2022. Association of the Gut Microbiota With Cognitive Function in Midlife. JAMA Netw Open. 5(2):e2143941.

Minter MR, Hinterleitner R, Meisel M, Zhang C, Leone V, Zhang X, Oyler-Castrillo P, Zhang X, Musch MW, Shen X et al. 2017. Antibiotic-induced perturbations in microbial diversity during post-natal development alters amyloid pathology in an aged APPSWE/PS1DeltaE9 murine model of Alzheimer’s disease. Sci Rep. 7(1):10411.

Minter MR, Zhang C, Leone V, Ringus DL, Zhang X, Oyler-Castrillo P, Musch MW, Liao F, Ward JF, Holtzman DM et al. 2016. Antibiotic-induced perturbations in gut microbial diversity influences neuro-inflammation and amyloidosis in a murine model of Alzheimer’s disease. Sci Rep. 6:30028.

Nakae S, Komiyama Y, Nambu A, Sudo K, Iwase M, Homma I, Sekikawa K, Asano M, Iwakura Y. 2002. Antigen-specific T cell sensitization is impaired in IL-17-deficient mice, causing suppression of allergic cellular and humoral responses. Immunity. 17(3):375–387.

O’Neill LA, Golenbock D, Bowie AG. 2013. The history of Toll-like receptors - redefining innate immunity. Nat Rev Immunol. 13(6):453–460.

Quan W, Luo Q, Hao W, Tomic I, Furihata T, Schulz-Schaffer W, Menger MD, Fassbender K, Liu Y. 2021. Haploinsufficiency of microglial MyD88 ameliorates Alzheimer’s pathology and vascular disorders in APP/PS1-transgenic mice. Glia. 69(8):1987–2005.

Radde R, Bolmont T, Kaeser SA, Coomaraswamy J, Lindau D, Stoltze L, Calhoun ME, Jaggi F, Wolburg H, Gengler S et al. 2006. Abeta42-driven cerebral amyloidosis in transgenic mice reveals early and robust pathology. EMBO Rep. 7(9):940–946.

Roberts KF, Elbert DL, Kasten TP, Patterson BW, Sigurdson WC, Connors RE, Ovod V, Munsell LY, Mawuenyega KG, Miller-Thomas MM et al. 2014. Amyloid-beta efflux from the central nervous system into the plasma. Ann Neurol. 76(6):837–844.

Shinohara M, Tachibana M, Kanekiyo T, Bu G. 2017. Role of LRP1 in the pathogenesis of Alzheimer’s disease: evidence from clinical and preclinical studies. J Lipid Res. 58(7):1267–1281.

Storck SE, Meister S, Nahrath J, Meissner JN, Schubert N, Di Spiezio A, Baches S, Vandenbroucke RE, Bouter Y, Prikulis I et al. 2016. Endothelial LRP1 transports amyloid-beta(1-42) across the blood-brain barrier. J Clin Invest. 126(1):123–136.

Vogt NM, Kerby RL, Dill-McFarland KA, Harding SJ, Merluzzi AP, Johnson SC, Carlsson CM, Asthana S, Zetterberg H, Blennow K et al. 2017. Gut microbiome alterations in Alzheimer’s disease. Sci Rep. 7(1):13537.

Xie J, Bruggeman A, De Nolf C, Vandendriessche C, Van Imschoot G, Van Wonterghem E, Vereecke L, Vandenbroucke RE. 2023. Gut microbiota regulates blood-cerebrospinal fluid barrier function and Abeta pathology. EMBO J. 42(17):e111515.

Xu S, Cao X. 2010. Interleukin-17 and its expanding biological functions. Cellular & Molecular Immunology. 7(3):164–174.

Zeng X, Li J, Shan W, Lai Z, Zuo Z. 2023. Gut microbiota of old mice worsens neurological outcome after brain ischemia via increased valeric acid and IL-17 in the blood. Microbiome. 11(1):204.

Zhan X, Stamova B, Jin LW, DeCarli C, Phinney B, Sharp FR. 2016. Gram-negative bacterial molecules associate with Alzheimer disease pathology. Neurology. 87(22):2324–2332.

Zhang T, Gao G, Kwok LY, Sun Z. 2023. Gut microbiome-targeted therapies for Alzheimer’s disease. Gut Microbes. 15(2):2271613.

Zhang Y, Shen Y, Liufu N, Liu L, Li W, Shi Z, Zheng H, Mei X, Chen CY, Jiang Z et al. 2023. Transmission of Alzheimer’s disease-associated microbiota dysbiosis and its impact on cognitive function: evidence from mice and patients. Mol Psychiatry. doi: 10.1038/s41380-023-02216-7.

Zhao Y, Jaber V, Lukiw WJ. 2017. Secretory Products of the Human GI Tract Microbiome and Their Potential Impact on Alzheimer’s Disease (AD): Detection of Lipopolysaccharide (LPS) in AD Hippocampus. Front Cell Infect Microbiol. 7:318.

Zhou Y, Xie L, Schroder J, Schuster IS, Nakai M, Sun G, Sun YBY, Marino E, Degli-Esposti MA, Marques FZ et al. 2023. Dietary Fiber and Microbiota Metabolite Receptors Enhance Cognition and Alleviate Disease in the 5xFAD Mouse Model of Alzheimer’s Disease. J Neurosci. 43(37):6460–6475.

