## Supplementary figures and images for "Modulation of Alzheimer’s Disease Brain Pathology in Mice by Gut Bacterial Deletion: The Role of Il-17a and Microglial MyD88"

### Supplementary Figure 1, Antibiotic treatment dramatically changes the bacterial composition in the gut of APP-transgenic mice.

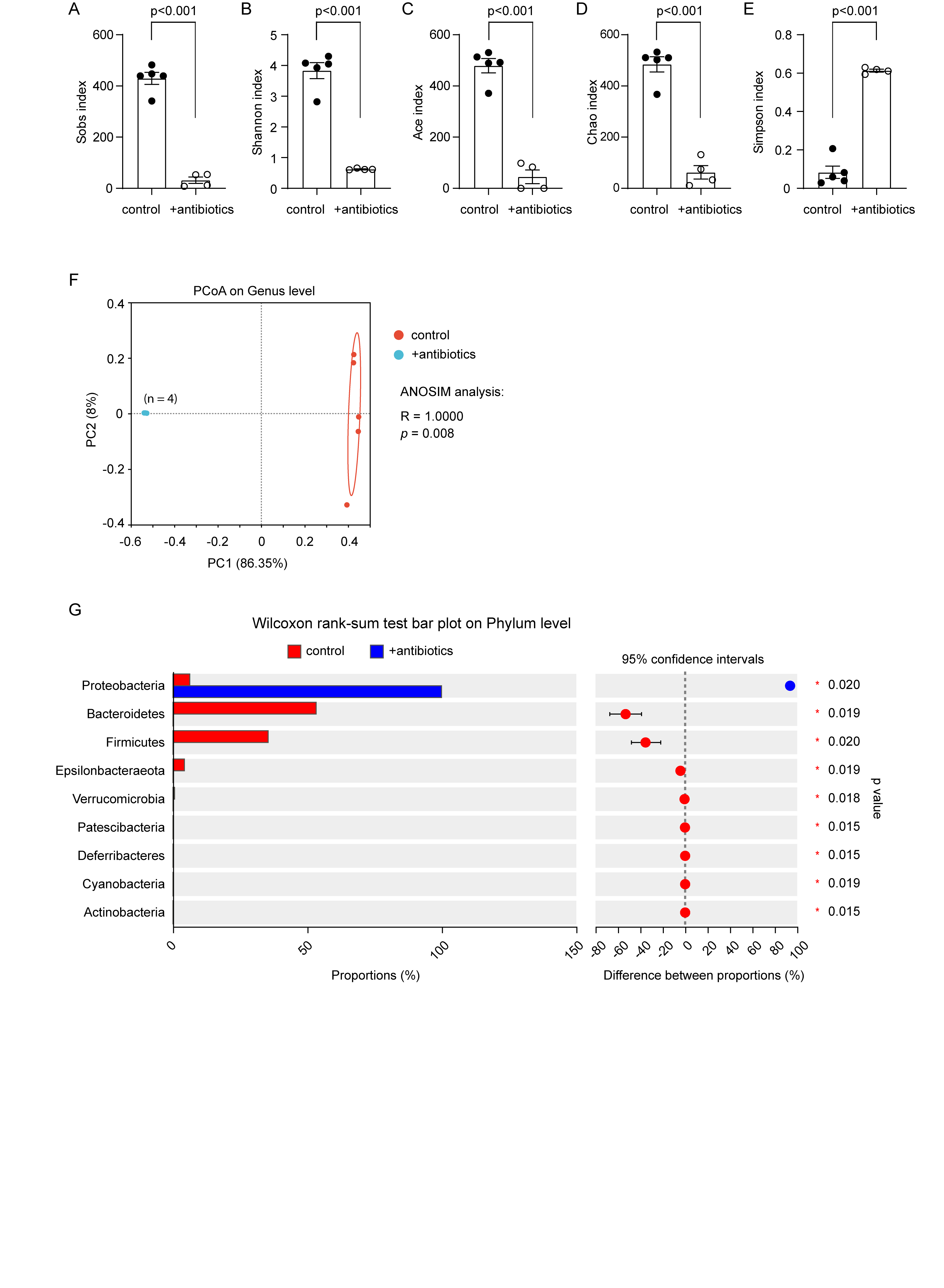
